# tensorOmics: Data integration for longitudinal omics data using tensor factorisation

**DOI:** 10.64898/2026.02.10.705179

**Authors:** Saritha Kodikara, Brendan Lu, Shuhe Wang, Ziqi Lei, Kim-Anh Lê Cao

## Abstract

Longitudinal multi-omics studies track molecular changes across biological layers and over time, but repeated measurements from the same subject introduce dependencies that must be accounted for. Conventional matrix-based approaches often flatten the data, obscuring this temporal structure. Tensor methods preserve the multi-way organisation; however, existing approaches are either unsupervised or limited to a single omic, leaving few options for phenotype-guided analysis across multiple longitudinal omics datasets.

We present tensorOmics, a framework comprising five tensor methods including an R implementation of TCAM as tensor PCA, spanning exploratory analysis, regression, classification and multi-omics integration. By representing each subject’s trajectory in an orthogonal temporal basis, tensorOmics retains subjects as the unit of analysis and enables fitted models to score new subjects. We evaluated the framework using simulations with known temporal signals and in three large public longitudinal cohorts. These evaluations assessed the recovery of group differences encoded in trajectory shape, prediction in held-out subjects and multi-omics integration. Across three biological case studies, our analyses identified relevant temporal patterns and molecular features. Together, these capabilities make tensorOmics a unified framework for analysing longitudinal single- and multi-omic data while preserving their temporal structure.

**Author summary:** Longitudinal multi-omics studies measure several types of molecules repeatedly in the same subjects or experimental units. Each omics dataset is naturally a three-dimensional array, yet standard methods often reshape it into a matrix, which can obscure temporal ordering and the dependence among repeated measurements. We developed tensorOmics to analyse longitudinal omics data in their natural form: subjects × molecular features × time. The framework includes exploratory and supervised methods for analysing subjects measured in multiple omics datasets, while treating each subject’s complete trajectory as one observation. It identifies temporal patterns and the molecular features that contribute to them, relates patterns across omics layers, classifies subjects into groups and scores new subjects.

We evaluated tensorOmics using simulations with known temporal patterns and subject-level prediction in three large public longitudinal cohorts. The benchmarks spanned controlled temporal signals and real-world variation in both single- and multi-omic settings. Three case studies further demonstrated how to interpret the identified temporal patterns and their contributing molecular features. tensorOmics is available as an R package for researchers studying how molecular systems change over time.

## 1 Introduction

Longitudinal multi-omics experiments are increasingly used to investigate disease progression, treatment response and biomarker discovery (Wang and Agapito, 2025, Mishra et al., 2022, Park et al., 2023, Ahmed et al., 2024). By monitoring subject-specific molecular changes over time, these studies can reveal coordinated shifts across biological layers that are missed by static cross-sectional designs. They can also distinguish temporal changes within subjects from variation between subjects, helping to identify molecular signatures of individual disease trajectories and treatment responses. This temporal perspective is important for understanding adaptive biological processes and for predicting clinical outcomes from early molecular changes.

Despite the increasing availability of longitudinal multi-omics datasets, analytical tools that both preserve their multi-way structure and incorporate clinical information remain limited (Baião et al., 2025, Luo et al., 2024). The high dimensionality of each omics layer increases further when several layers are integrated over time, while variation between subjects can obscure temporal patterns shared across subjects. Missing follow-up time points lead to irregular sampling and complicate trajectory reconstruction. Differences in the statistical properties of omics data pose an additional challenge: microbiome data, for example, are sparse and compositional, whereas proteomic measurements often follow log-normal distributions. A unified framework must therefore account for these differences while detecting group effects expressed through trajectory shape rather than mean abundance. However, such effects may be obscured by large, stable between-subject differences that dominate the variance.

Most methods for multi-omics integration were developed for cross-sectional data and can be broadly grouped into unsupervised and supervised approaches. Unsupervised methods uncover structure without prior knowledge of phenotypes or sample labels, either identifying new phenotypic groups or capturing sources of variation shared across omics layers. Similarity Network Fusion (Wang et al., 2014) and Bayesian Consensus Clustering (Kirk et al., 2012), for example, identify phenotypic subgroups, whereas joint Non-negative Matrix Factorization, Joint and Individual Variation Explained (JIVE) (Lock et al., 2013), sparse MultiBlock Partial Least Squares (Li et al., 2012), sparse generalized canonical correlation analysis (sGCCA) (Tenenhaus and Tenenhaus, 2011, Tenenhaus et al., 2014), and Multi-Omics Factor Analysis (MOFA) (Argelaguet et al., 2018) capture shared structure. Supervised methods instead relate omics data to known subject labels or phenotypes, either categorical (classification) or continuous (regression), to identify molecular features that discriminate between groups or predict outcomes. Data Integration Analysis for Biomarker discovery using Latent cOmponents (DIABLO), for example, identifies multi-omic biomarkers that link molecular patterns across biological layers to known phenotypes (Singh et al., 2019). To analyse longitudinal data, however, these matrix-based methods must flatten the three-way array, commonly by assigning one row to each subject–time point or one column to each feature–time point. Neither arrangement explicitly models temporal order; the former treats repeated measurements from the same subject as independent observations, and the latter inflates the number of features by the number of time points.

Tensors extend matrices beyond two dimensions and naturally organise longitudinal omics measurements as subjects × omics features × time points. This representation preserves the multi-way structure and enables subjects, molecular features and temporal dynamics to be modelled jointly. Several unsupervised tensor methods have been developed for longitudinal single-omic data, including Context-aware Tensor Factorization (CTF) (Martino et al., 2021), microTensor (Ma and Li, 2023), TCAM (Mor et al., 2022), FTSVD (Han et al., 2024), EMBED (Shahin et al., 2023), and TEMPTED (Shi et al., 2024). These methods use tensor decomposition to identify low-dimensional structure that captures temporal variation, but represent time differently. For example, TCAM uses a fixed orthogonal basis, TEMPTED estimates a smooth function of continuous time, and PARAFAC (Harshman et al., 1970), a classical multi-way decomposition, estimates a free temporal loading vector for each component. Their focus on unsupervised, single-omic analysis leaves phenotype-guided analysis across multiple longitudinal omics blocks unresolved.

Several supervised, design-based and multi-view alternatives address individual aspects of this problem. N-PLS (Bro, 1996) extends partial least squares (PLS) to three-way arrays and relates one block to a response through a freely estimated weight vector over time. Repeated measures ANOVA simultaneous component analysis+ (RM-ASCA+) (Madssen et al., 2021) tests the group-by-time interaction at the population level, but is not designed to classify individual subjects. MEFISTO (Velten et al., 2022) integrates several data views using a Gaussian-process prior on time, but is neither tensor-based nor supervised. Consequently, no existing method provides supervised analysis (classification or regression) across several longitudinal omics blocks.

We introduce tensorOmics, a framework of five tensor methods for dimensionality reduction and the integration of longitudinal multi-omics data. The framework combines tensor decomposition with regularised generalised canonical correlation analysis (RGCCA) (Tenenhaus and Tenenhaus, 2011), supports unsupervised and supervised analyses, and accommodates any number of data blocks. Its five methods target distinct analytical objectives: tensor PCA for single-omic exploratory analysis, tensor partial least squares for relating pairs of omics datasets or modelling continuous outcomes, tensor partial least squares discriminant analysis for single-omic classification, block tensor PLS for multi-omics integration, and block tensor PLS-DA for classification using multiple omics datasets.

Within tensorOmics, tensor PCA provides the unsupervised single-omic foundation through an R implementation of TCAM (Mor et al., 2022), while the remaining four methods extend this representation to regression, classification and multiblock integration. All five methods represent time using a fixed orthogonal basis, the discrete cosine transform, and construct each component from a single basis term. For classification, the centred class-indicator matrix is applied unchanged across all temporal faces, and a scaled score selects the face, preventing stable differences between subjects from masking differences in trajectory shape. Fitted models can then be evaluated on held-out subjects.

We evaluated the methods through simulations with known informative features and temporal patterns. Using these simulations, we assessed the recovery of a group difference from trajectory shape, the role of face scaling when between-subject variation is large, the influence of feature scaling when comparing temporal representations, and the effect of disagreement among omics layers in the timing of group differences. In three public multi-omics cohorts, we compared subject-level cross-validation across matrix and tensor methods, with a feature-scaling sensitivity analysis for single-block methods; multiblock comparisons were not matched for scaling. Finally, three case studies spanned different experimental designs and small sample sizes commonly encountered in longitudinal omics research and illustrated how components and loadings can be interpreted under realistic conditions.

## 2 Materials and methods

### 2.1 Data representation and temporal transformation

Our framework tensorOmics represents each longitudinal omic block as a tensor with dimensions subjects *×* omic features *×* time. It extends the single-block tensor PCA (tPCA) implemented by TCAM (Mor et al., 2022) to tensor partial least squares (tPLS), discriminant analysis (tPLS-DA), and their multiblock counterparts, block tPLS and block tPLS-DA.

#### 2.1.1 Notation and tensor operations

Tensors, matrices, vectors and scalars are denoted by bold italic uppercase, bold uppercase, underlined lowercase and lowercase symbols, respectively. A tensor is a multidimensional array, and its order is the number of dimensions, or modes. A third-order tensor ***A*** ∈ ℝ^*n×p×t*^ has three modes and can be viewed as a collection of *t* matrices of size *n× p*. Its element *A*_*i,j,k*_ is the observation for subject *i*, feature *j* and time *k*. Thus, mode 1 indexes subjects, mode 2 indexes features and mode 3 indexes time (Fig. 1A).

**Fig 1.**
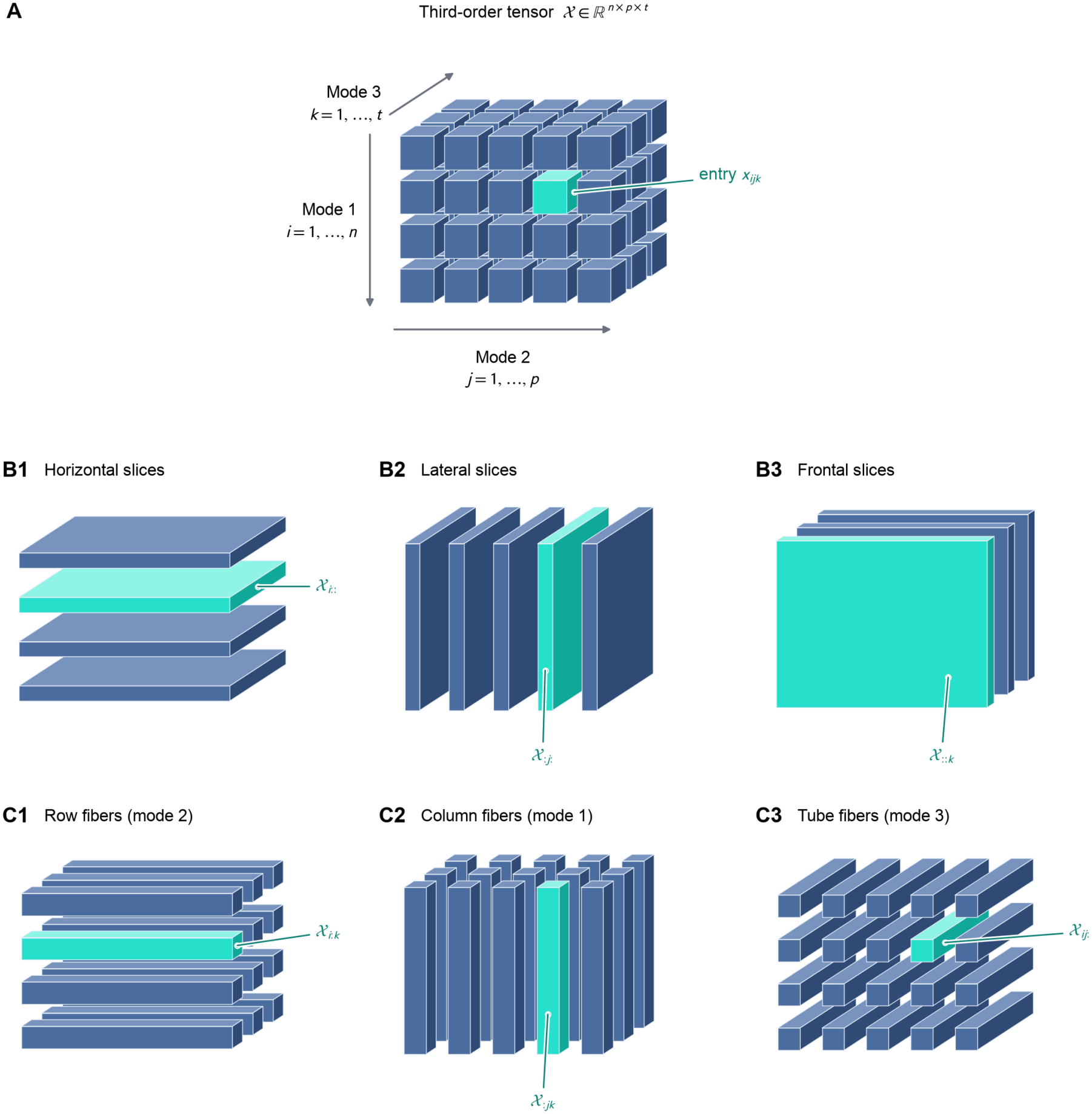
Structure of a third-order tensor and its slices and fibres. **A**: A third-order tensor ***A***∈ ℝ^*n×p×t*^, where *A*_*i,j,k*_ is the observation for subject *i*, feature *j* and time *k*. The modes correspond to the tensor dimensions: mode 1 (*i*) indexes subjects, mode 2 (*j*) indexes features and mode 3 (*k*) indexes time. **B1–B3**: Matrix-valued slices obtained by holding one mode constant: a horizontal slice (mode 1), lateral slice (mode 2) and frontal slice (mode 3). **C1–C3**: Vector-valued fibres obtained by holding two modes constant: a row fibre varies along mode 2, a column fibre along mode 1 and a tube fibre along mode 3.

A slice is obtained by fixing one index and is therefore a matrix. Fixing the subject, feature or time index gives a horizontal slice ***A***_*i*,:,:_, lateral slice ***A***_:,*j*,:_ or frontal slice ***A***_:,:,*k*_, respectively (Fig. 1B1–B3) (Kilmer et al., 2021). Frontal slices are also called faces, and face-wise operations act on each face separately. For example, the transpose ***A***^*⊤*^ ∈ ℝ^*p×n×t*^ is the face-wise transpose, (***A***^*⊤*^)_:,:,*k*_ = (***A***_:,:,*k*_)^*⊤*^. A fibre is a vector obtained by fixing two indices. The row, column and tube fibres are ***A***_*i*,:,*k*_, ***A***_:,*j,k*_ and ***A***_*i,j*,:_, respectively (Fig. 1C1–C3).

The face-wise product of 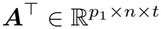 and 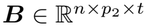multiplies their matching frontal matrices:

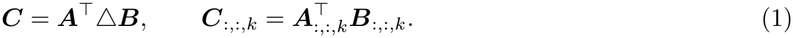

Here, *△* denotes the face-wise product and 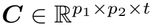, as illustrated in Fig. 2A.

**Fig 2.**
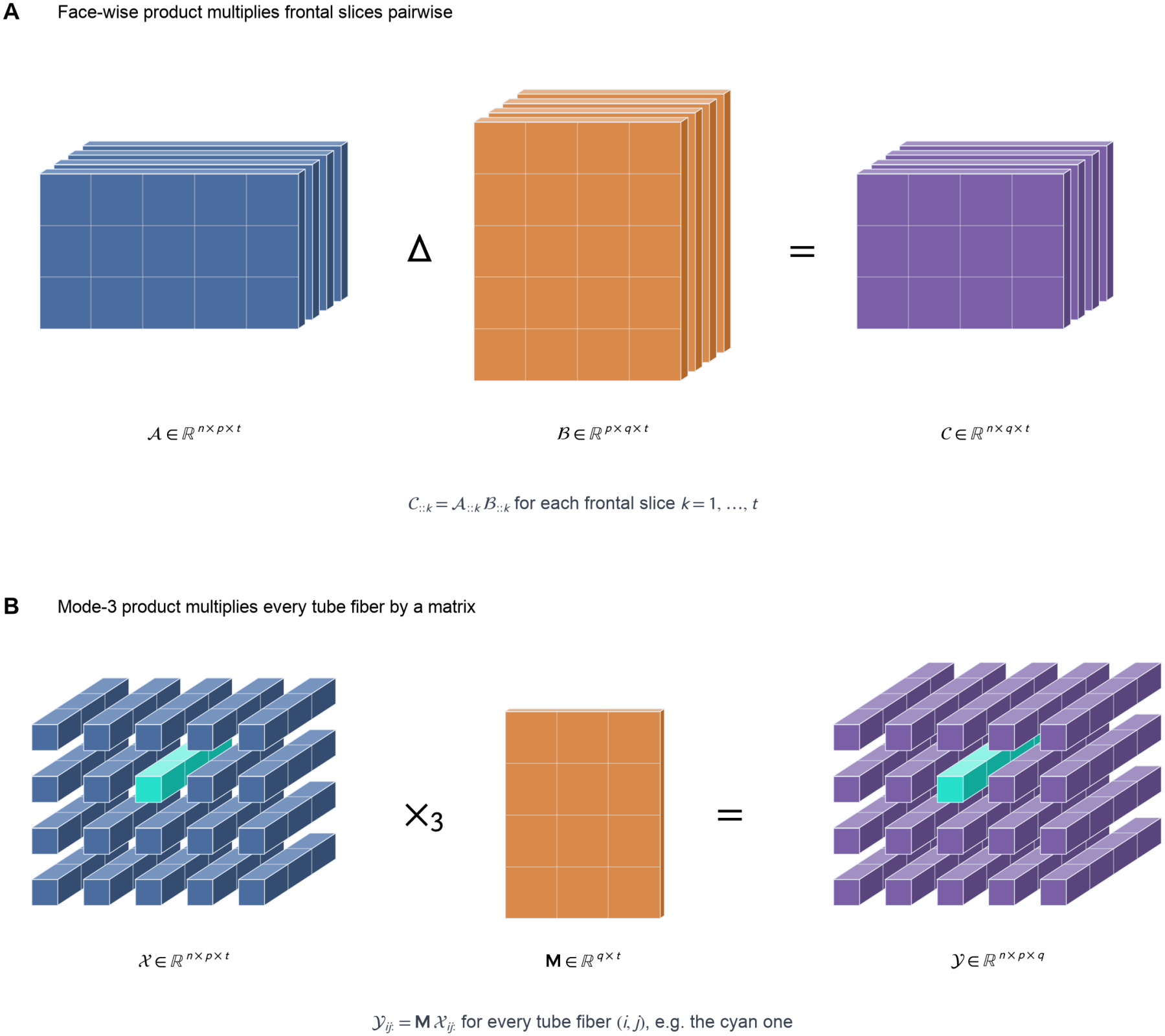
Tensor operations used by tensorOmics. **A**: In the face-wise product, 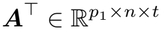 and 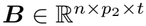 are combined by multiplying matching frontal slices. For each *k*, 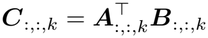, giving 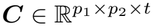. **B**: In the mode-3 product, ***A*** ℝ^*n×p×t*^ is multiplied along its third mode by the temporal-basis matrix **M** ∈ ℝ^*t×t*^. Each tube fibre is updated as ***Â***_*i,j*,:_ = **M*A***_*i,j*,:_. The matrix is square here because tensorOmics uses an orthonormal transform over the *t* time points.

The mode-3 product applies a matrix to every tube fibre. For ***X*** ∈ ℝ^*n×p×t*^ and **M** ∈ ℝ^*s×t*^,

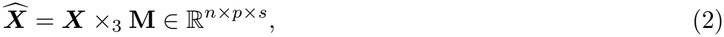

where *×*_3_ indicates multiplication along mode-3 (Fig. 2**B**). Specifically, each length-*t* tube fibre ***X***_*i,j*,:_ is mapped to the length-*s* fibre 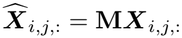. In tensorOmics, *s* = *t* and **M** is an orthonormal temporal-basis matrix, so **M** is square and **M**^*−*1^ = **M**^*⊤*^.

#### 2.1.2 Data representation and pre-processing

Let 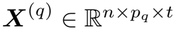 denote omic block *q*, where all *Q* blocks share subjects and an aligned, complete time grid. A longitudinal continuous response is a tensor ***Y***∈ ℝ^*n×m×t*^. A subject-level continuous response is an *n × m* matrix, and a categorical response is the centred indicator matrix **Y**_*c*_ ∈ ℝ^*n×m*^. Subject-level responses are not transformed over time, and the same matrix is supplied to every temporal face.

Each predictor tensor, and any longitudinal response tensor, is centred across subjects within every feature-time cell:

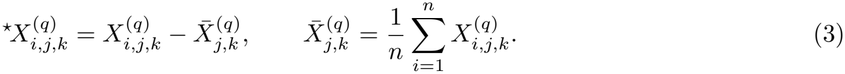

Subject-level responses are centred column-wise. For prediction and cross-validation, we use means estimated from the training subjects only. In tensorOmics, we do not standardise features or scale blocks, as these remain pre-processing choices (this is distinct from the face scaling of Eq. 8, which only affects face selection).

Note, however, that because the objectives maximise covariance, features or blocks with greater post-transformation variance can contribute more strongly.

#### 2.1.3 Temporal basis

Following TCAM Mor et al. (2022), our framework uses the orthonormal type-II discrete cosine transform (DCT-II). For time or visit positions *k, l* = 1, ߪ, *t*,

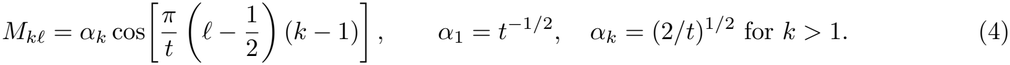

The transformed tensor is

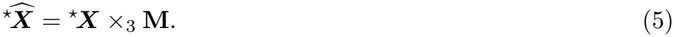

Face 1 is proportional to the average over time, while subsequent faces represent progressively higher-frequency contrasts (face 2 is a monotone trend, face 3 a U-shaped contrast up to sign, and so on). We retain all faces, but each component is constructed from one selected face, so its temporal pattern is the corresponding cosine basis function. The DCT uses ordered time positions and hence does not explicitly model unequal time intervals; faces describe patterns across visit order, not calendar time.

### 2.2 Component extraction and multiblock integration

tensorOmics combines transform-domain tensor decomposition (Kernfeld et al., 2015) with regularised generalised canonical correlation analysis (RGCCA) (Tenenhaus and Tenenhaus, 2011). A canonical analysis relates two or more omic blocks symmetrically, without designating any block as the response. A regression analysis instead designates a continuous or categorical response and estimates predictor components in relation to that response (as described in more detail in the Deflation paragraph). Thus, tPLS-DA and block tPLS-DA perform classification but use the response-directed PLS formulation and its regression deflation, with class membership represented by an indicator matrix.

Components are extracted from selected transformed faces and sequentially deflated (as defined below). Multiblock weights are estimated by NIPALS-type RGCCA iterations. In short, after centring and the DCT each face is an ordinary *n × p*_*q*_ matrix; every component is obtained by choosing one face *k*_*h*_ and then finding unit-norm feature weights on that face by a PLS-type (covariance-based) analysis.

#### Pairwise transform-domain decomposition

For two centred tensors, let 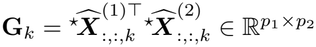.The t-SVDM2 operation applies an ordinary singular value decomposition to every face,

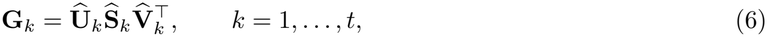

where the diagonal entries of **Ŝ**_*k*_ are the singular values. In Algorithm 1, the two centred tensors are transformed along mode 3, the face-wise covariance matrices are decomposed, and the resulting factor tensors are mapped back to the time domain. With identical tensors, Eq. 6 reduces to the face-wise decomposition used by t-SVDM.

##### Algorithm 1

t-SVDM2 decomposition of a face-wise covariance tensor

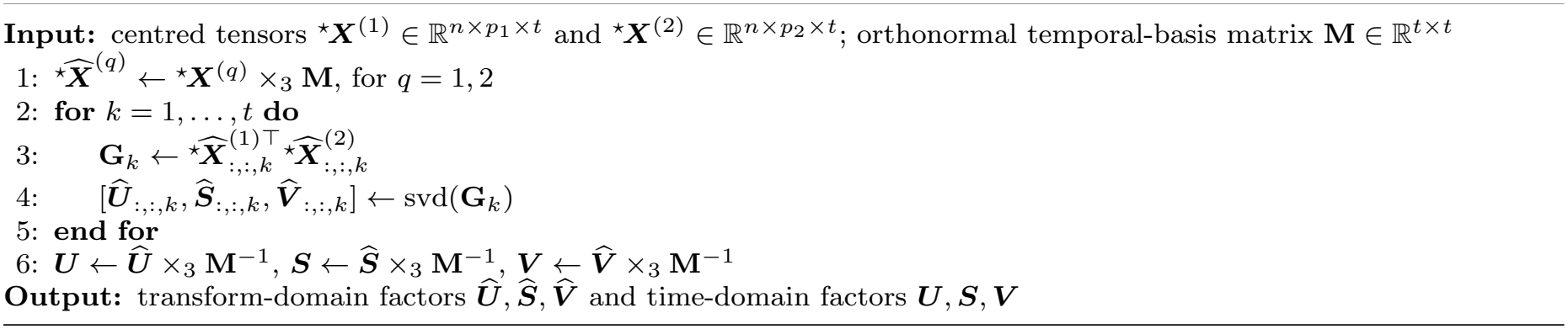

#### Component and face selection

For tPCA, an SVD is computed for every transformed face, and the singular values are ranked jointly across all singular-vector–face pairs. The *H* largest pairs define the retained components, their feature loadings and their selected faces (Mor et al., 2022).

For methods with two or more omic datasets, and optionally with a response, let 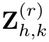 be the residual matrix for block *r* at face *k* before component *h*. We assume a subject-level response is shared across faces. The symmetric design matrix **C** = (*c*_*rs*_) has *c*_*rr*_ = 0 and non-negative off-diagonal entries. Each off-diagonal entry is a specified design weight ∈ [0, 1]: *c*_*rs*_ = 0 means that blocks *r* and *s* are not directly linked, whereas *c*_*rs*_ = 1 gives the pair full weight in the objective, so that larger values place more emphasis on agreement between the two blocks. With *B* blocks (*B* = *Q* without a response; *B* = *Q* + 1 when the response is appended as block *Q* + 1, Eq. 17), face *k* is scored by

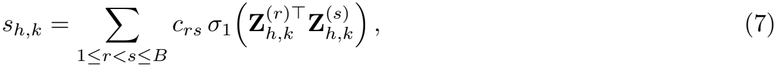

where *σ*_1_(·) is the largest singular value, so *s*_*h,k*_ is the strongest weighted cross-covariance attainable on face *k*. To compare faces on a common scale, supervised analyses use

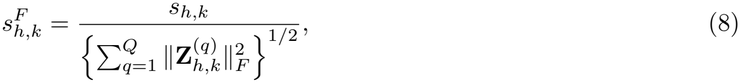

where the denominator contains the *Q* predictor blocks but not the response. The selected face is 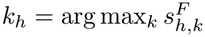 for supervised analyses and *k*_*h*_ = arg max_*k*_ *s*_*h,k*_ otherwise. Stable between-subject differences fall on the constant face 1 and inflate its covariance, so unscaled selection could favour face 1 even when a later face separates the groups better. The scaling, analogous to rescaling a covariance into a correlation, compares faces on a common scale. Scaling affects face selection only; weights, scores and deflation all use the unscaled matrices.

At the selected face, tensor RGCCA solves

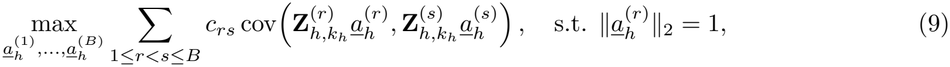

using the Horst scheme (identity scheme). Because the weights are constrained to unit Euclidean norm, rather than the scores being standardised to unit variance, the criterion is covariance-based (PLS-type, shrinkage parameter *τ* = 1 in RGCCA), so “canonical” refers to symmetric deflation, not to a correlation criterion. We define the score for block *r* as 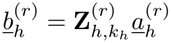.

#### Deflation

Deflation is the sequential removal, after each component, of the rank-one variation represented by its scores and loadings, so that the next component is fitted to the remaining variation. Canonical and regression deflation differ in how we calculate the residuals from the blocks and response.

In canonical analyses (symmetric deflation), every block is deflated by its own score. For block *r*,

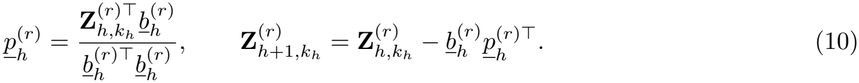

Only face *k*_*h*_ is updated and all other faces are carried forward unchanged. In regression analyses (asymmetric deflation), the predictor is deflated as above and a subject-level response residual **Y**_*h*_ is deflated by the predictor score:

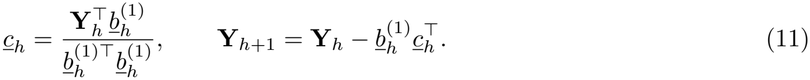

With *Q* predictors, the block-specific response coefficient is 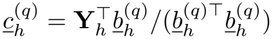, and the shared response is deflated by the average contribution,

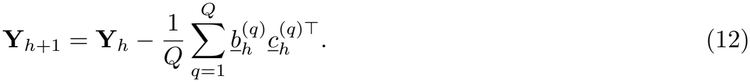

The preceding updates apply to a subject-level response, which has no time mode. A longitudinal response is a tensor with a time mode: its residual is transformed by **M**, updated on the face selected for the current component using the same score-based deflation, and then mapped back to the time domain with **M**^*−*1^. The resulting tensor is the response residual used for the next component. Before any deflation, canonical and regression configurations optimise the same covariance criterion and therefore yield the same first component for a given pair of blocks. Later components can differ because canonical deflation updates each block using its own score, whereas regression deflation updates the response using the predictor score or the average contribution of the predictor-block scores (Wegelin et al., 2000, Rohart et al., 2017). Algorithm 2 summarises these updates within the sequential component-extraction procedure; the extended pseudo-algorithm is provided in Supplemental Algorithm A.

##### Algorithm 2

Component extraction for tPLS and block.tPLS methods

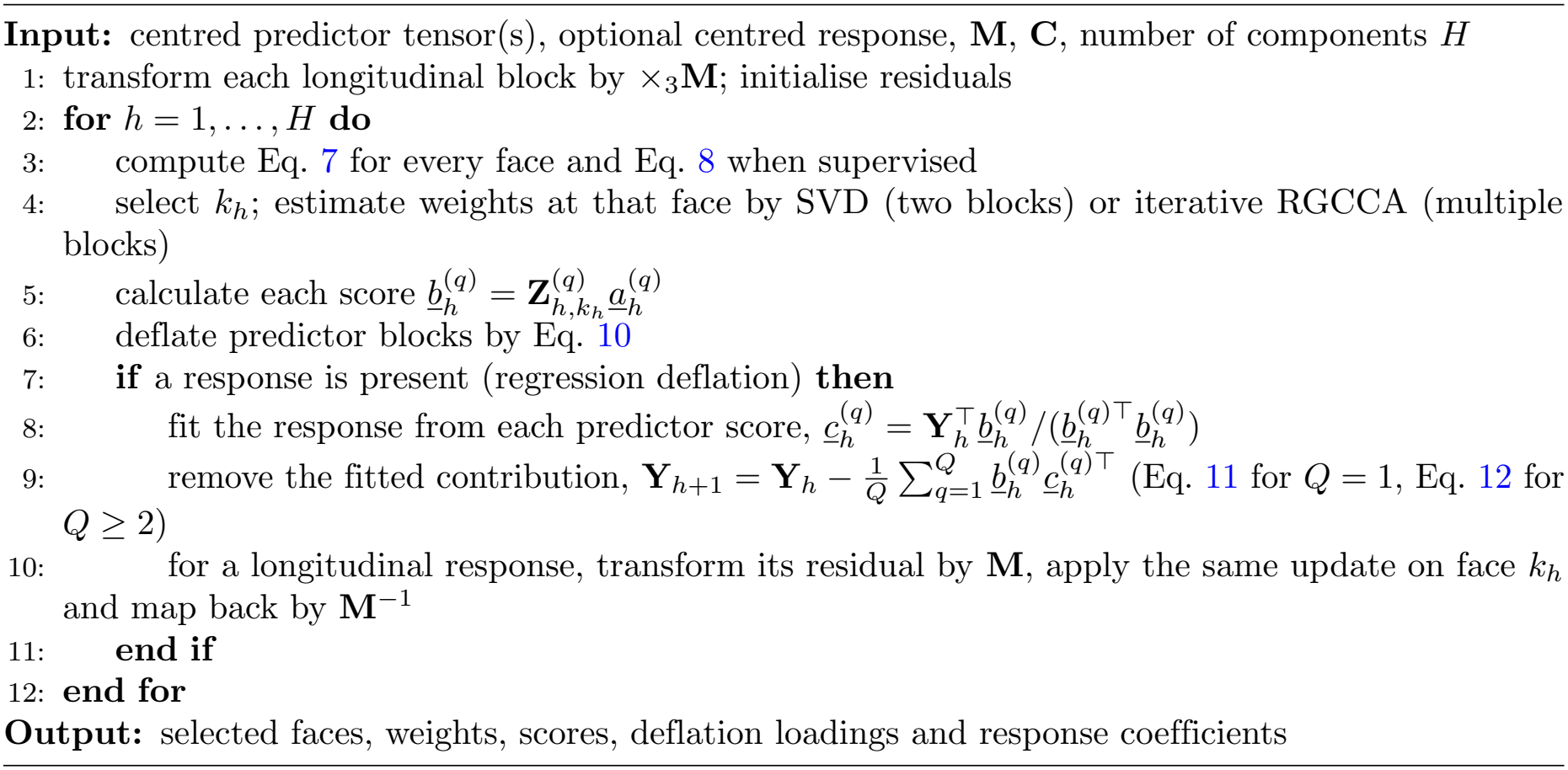

### 2.3 Method configurations

The five method families form seven configurations according to the number of predictor blocks, the presence of a response and its type (Fig. 3; Table 1). tPCA uses within-block SVD, two-block tPLS uses t-SVDM2, and multiblock methods use iterative RGCCA.

**Table 1.** tensorOmics method configurations and their objectives. *Q* is the number of omic predictor blocks; the response is not included in *Q*. A continuous response may be a subject-level matrix or a longitudinal tensor, for example a second omic block.

| Method and configuration | Omic predictors | Designated response | Deflation | Objective |
|---|---|---|---|---|
| tPCA | 1 | none | not required (SVD) | Eq. 13: within-block variance |
| tPLS (canonical) | 2 | none | canonical | Eq. 14: covariance between omics |
| tPLS (regression) | 1 | continuous | regression | Eq. 15: predictor–response covariance |
| tPLS-DA | 1 | categorical | regression | Eq. 15: predictor–class covariance |
| block.tPLS (canonical) | *Q* ≥ 3 | none | canonical | Eq. 16: connected omic–omic covari-ance |
| block.tPLS (regression) | *Q* ≥ 2 | continuous | regression | Eq. 17: connected covariance with re-sponse |
| block.tPLS-DA | *Q* ≥ 2 | categorical | regression | Eq. 17: connected covariance with class |

**Fig 3.**
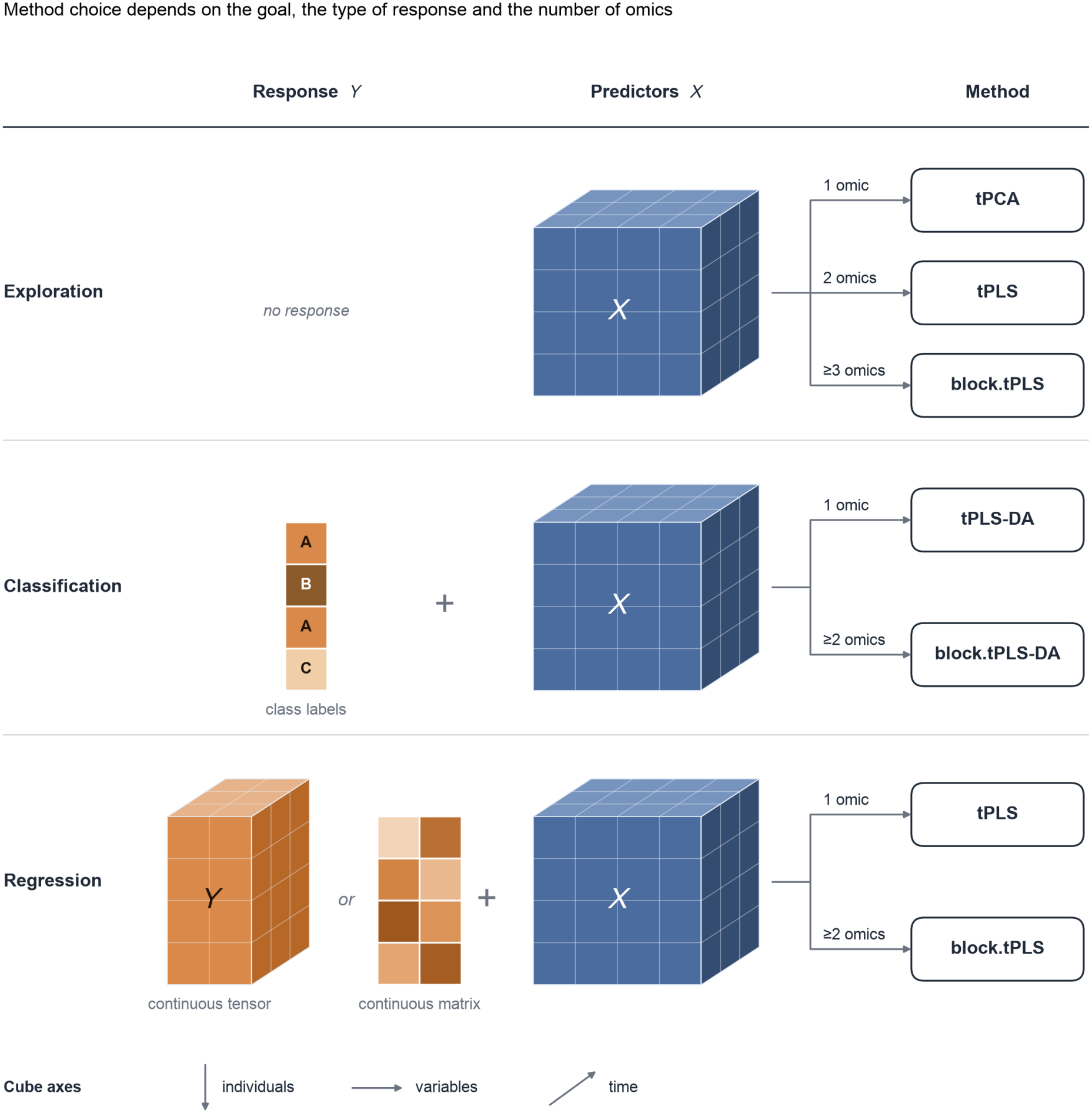
Method selection by analytical goal, response type and number of omic blocks. tPLS and block.tPLS use canonical deflation without a response and regression deflation with a response.

At component *h, Q* denotes the number of omic predictor blocks, *k*_*h*_ is the selected temporal face, and 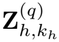 is the residual matrix for predictor block *q*. Its component score is 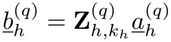. When a response is present, define its residual block at component *h* as

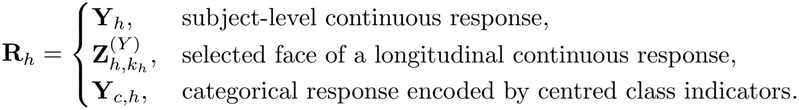

Thus, the response-directed objectives have the same mathematical form for regression and classification; the response encoding determines which method is fitted.

#### Single- and two-block objectives

tPCA uses one omic block and no response, and maximises within-block variance:

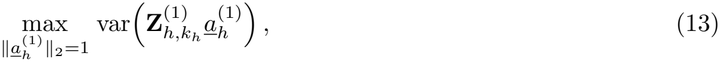

Canonical tPLS uses two omic blocks symmetrically, with neither designated as the response, and maximises their covariance:

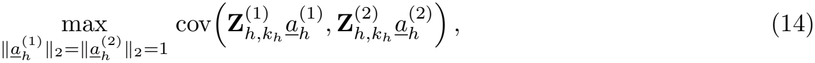

Response-directed tPLS uses one omic predictor and the response block **R**_*h*_:

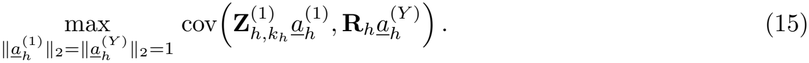

Equation 15 defines tPLS regression when **R**_*h*_ is continuous and tPLS-DA when **R**_*h*_ = **Y**_*c,h*_.

#### Multiblock objectives

Canonical block.tPLS has *Q* omic blocks and no response. It maximises the weighted covariance summed over the connected omic–omic pairs specified by the design matrix **C** = (*c*_*fg*_):

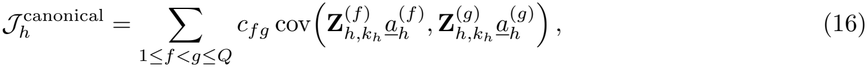

For response-directed block.tPLS, the response is appended as block *Q* + 1, with 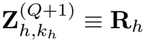. The objective then includes all connected omic–omic and omic–response pairs:

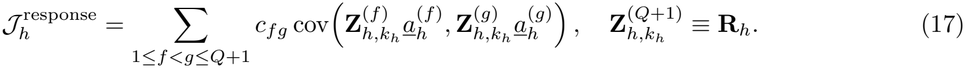

Equation 17 defines block.tPLS regression when **R**_*h*_ is continuous and block.tPLS-DA when **R**_*h*_ = **Y**_*c,h*_. All block weights have unit Euclidean norm. With two omic blocks and no response, Eq. 16 is the canonical tPLS objective; we therefore use the name canonical block.tPLS for three or more omic blocks. The distinction is computational: tPLS uses the direct t-SVDM2 solution, whereas block.tPLS uses iterative RGCCA.

#### 2.3.1 tPCA: tensor principal component analysis

tPCA is the single-omic unsupervised method. It ranks singular-vector–face pairs by explained variance (Eq. 13) and returns subject scores, feature loadings and the selected temporal faces.

tPCA is designed for exploratory analysis. Typical questions include: *What are the dominant temporal patterns in the dataset? Do subjects cluster by treatment or disease status? Which features drive these differences?* Ordination of subject scores reveals major trajectory patterns, while loadings highlight features such as taxa, transcripts, or metabolites contributing to these trends.

#### 2.3.2 tPLS: tensor partial least squares

tPLS maximises covariance between one predictor tensor and either a second omic or a continuous response (refer to Eq. 14 for a second omic, and to Eq. 15 for a response). Two omics without a designated response use canonical deflation; a designated response uses regression deflation. A longitudinal response is transformed as a tensor, whereas a subject-level response is shared across faces. The loadings describe the fitted covariance structure.

This method addresses questions such as: *Do cytokine and microbiome dynamics evolve together? (2-omics & no dedicated response → tPLS canonical deflation)*; *Can gene expression trajectories predict changes in protein expression? (1-omic & another omic as response → tPLS regression deflation)*; *Can gene expression trajectories predict changes in a specific clinical marker? (1-omic & continuous clinical marker → tPLS regression deflation)*. The analysis highlights subsets of features from each dataset that jointly explain the shared temporal variation, offering interpretable links between molecular and phenotypic layers or between two different molecular layers.

#### 2.3.3 tPLS-DA: tensor partial least squares discriminant analysis

##### Response construction and face scaling

tPLS-DA replaces the second block with the centred indicator matrix **Y**_*c*_ (Eq. 15). The objective is the covariance between the predictor score and the score of the class indicators. Since group membership is defined at the subject level, **Y**_*c*_ is supplied unchanged to every face and deflated once per component using Eq. 11. Faces are selected using Eq. 8. For one predictor, the denominator is 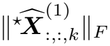. The scaling affects face selection only.

This approach is suited to questions such as: *Can subject trajectories be classified by disease subtype or treatment response? (1-omic & categorical response → tPLS-DA)* The results provide trajectory-aware molecular signatures that are predictive of outcome categories.

#### 2.3.4 block.tPLS: multiblock tensor partial least squares

block.tPLS extends tPLS to *Q ≥* 2 omic predictors (Eq. 16). Without a response it is canonical block.tPLS, which we define for *Q ≥* 3 because two omic blocks reduce to canonical tPLS; with a continuous response it applies for *Q ≥* 2 (Table 1). The design matrix sets the connected pairs and their weights. Predictors are deflated by their own scores; with a response, the response is deflated by the average block contribution (Eq. 12).

This approach is suited to questions such as: *What is the shared structure across microbiome, metabolome, and proteome trajectories? (3-omics & no dedicated response → block*.*tPLS canonical)*; *Can integrated microbiome, metabolome, and proteome trajectories predict disease progression? (3-omics & continuous clinical outcome as response → block*.*tPLS regression)*; *Can combined transcriptome and epigenome dynamics predict changes in a downstream omic layer? (2-omics & another omic as response → block*.*tPLS regression)*. The analysis highlights subsets of features from each dataset that jointly explain the shared temporal variation, offering interpretable links across multiple molecular layers or between multiple molecular layers and phenotypic outcome.

#### 2.3.5 block.tPLS-DA: multiblock tensor discriminant analysis

block.tPLS-DA adds **Y**_*c*_ as block *Q* + 1 (Eq. 17) and uses the scaled face criterion. Positive omic–omic design weights allow cross-omic agreement, as well as omic–outcome covariance, to influence the selected face.

This method is designed for questions such as: *Can combined transcriptome and epigenome dynamics distinguish responders from non-responders over time? (2-omic & categorical response → block*.*tPLS-DA)* The output is a trajectory-aware, integrative molecular signature that can be validated in independent cohorts.

### 2.4 Prediction and validation for new subjects

A fitted model projects a new subject without re-estimating the basis, faces or component directions. This requires a complete trajectory on the training time grid and does not forecast an unobserved time. For predictor block *q*, the new trajectory is centred by the training mean and transformed by the fitted basis:

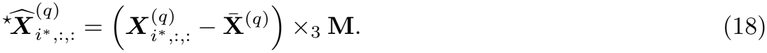

A tPCA score projects the selected face onto its retained right singular vector. For PLS methods, let 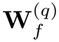 and 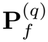 contain the weights and deflation loadings for components selected from face *f*. The face-specific rotation is

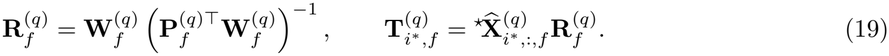

Components from the same face are projected jointly, preserving the fitted deflation.

Single-block classification uses nearest training-score centroids (Euclidean distance in the space of all *H* component scores). For multiblock classification, block scores are first averaged component-wise,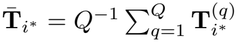, and the same training-centroid rule is applied to this consensus score vector.

For a subject-level continuous response, single- and multiblock predictions are

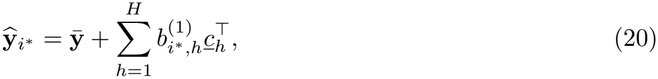

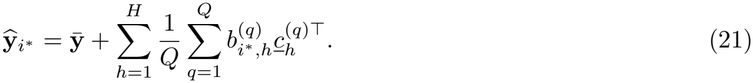

For a longitudinal response, contributions are accumulated on their selected faces and returned to the time domain with **M**^*−*1^. All prediction quantities, including means and class centroids, are estimated within each training fold.

### 2.5 Interpretation

#### Inputs and outputs

Tensor methods analyse longitudinal data as an array with dimensions subjects *×* molecular features *×* time and produce two complementary plots. A subject plot shows where each subject sits in a low-dimensional space defined by the components, and a loading plot shows which features contribute most to each component. To show how to read these plots, we use a small simulated dataset of 10 features measured in 10 subjects from two groups at five time points (Fig. 4**A**). We designed features V1–V5 to differ between the groups; V6–V10 contain no group signal.

**Fig 4.**
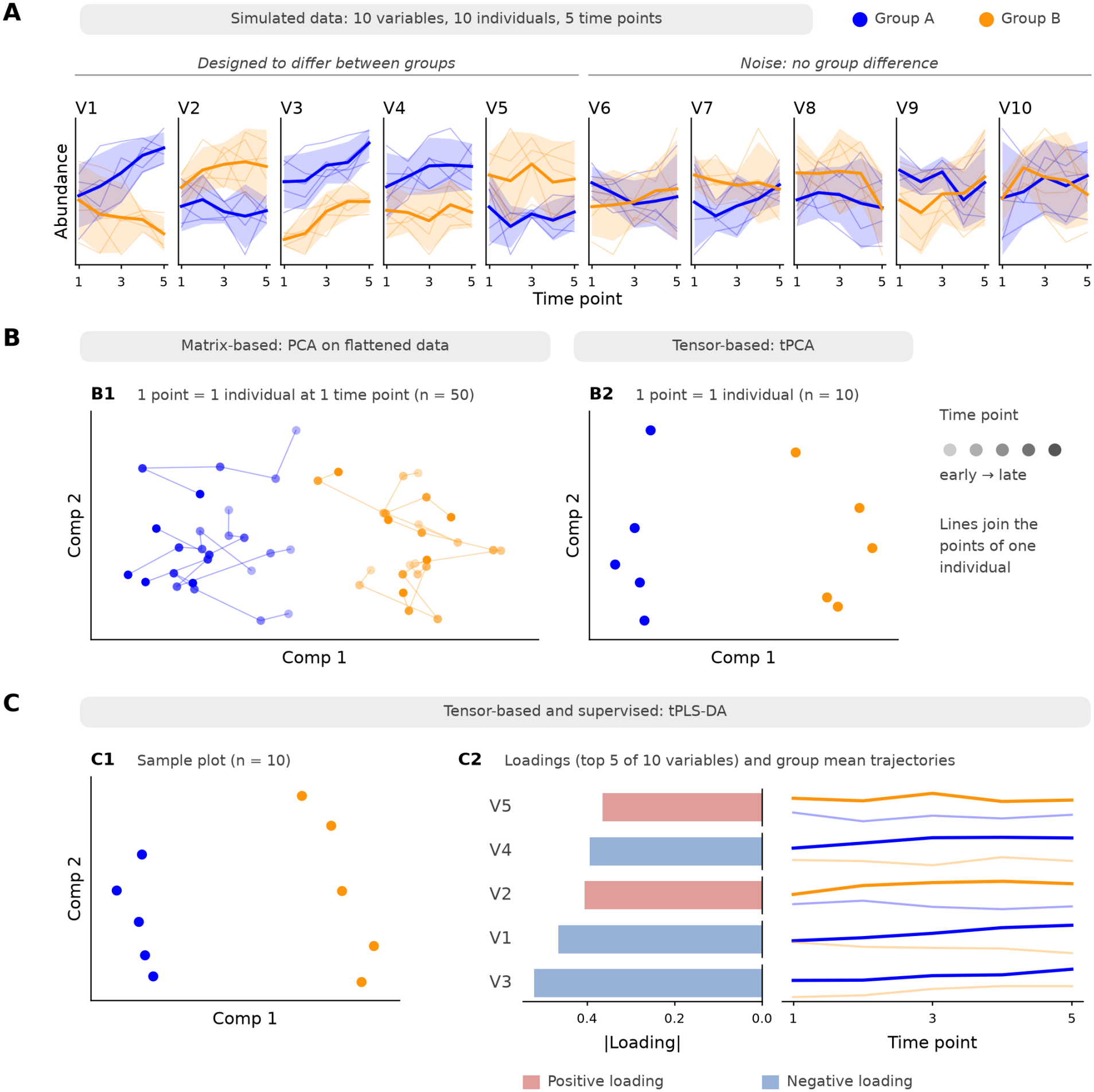
Interpretation of matrix and tensor outputs using simulated longitudinal data from 10 subjects, 10 features and five time points. **A**: subject and group mean trajectories. V1–V5 differ between groups, whereas V6–V10 are noise. **B**: Unsupervised sample and subject plots. **B1**: Principal component analysis (PCA) of the stacked subject-time matrix, with one point per subject and time. **B2**: Tensor principal component analysis (tPCA), with one point per subject trajectory. **C**: Tensor partial least squares discriminant analysis (tPLS-DA). **C1**: Fitted subject scores, showing separation along Component 1. **C2**: The five largest absolute Component 1 loadings and their corresponding group mean trajectories. All five are features containing the simulated group signal. Loading signs are relative to the subject plot orientation, and the curves describe group means. Component 2 completes the two-dimensional display but is not an additional discriminant direction in this two-group example.

#### Matrix-based versus tensor-based sample or subject plots

Fig. 4**B** shows how the two approaches treat repeated measurements. We refer to the matrix-based PCA plot as a *sample plot*, since each point represents one sample (a subject–time observation), and to the tensor-based tPCA plot as a *subject plot*, since each point is one subject. Standard PCA requires a matrix, so we stack the data with one row per subject-time combination. PCA treats each row as an independent observation. Each subject therefore appears as several points, one for each time point. tPCA instead keeps each subject as the unit of analysis and represents the subject’s complete trajectory by a single point. In this example, both methods separate the groups along Component 1. We show the comparison to illustrate how each method represents repeated measurements, not to compare their performance.

#### Supervised subject plot

In the tPLS-DA subject plot (Fig. 4**C1**) where each point represents one subject’s complete trajectory, groups A and B separate along Component 1. This graphical representation shows that the fitted model has captured the simulated group difference (the group labels have been included in this supervised analysis). The plot therefore describes the fitted model; it does not show how well the model predicts held-out subjects, which we evaluate in the benchmarks (Section 2.6).

#### Loading plot

The loading plot shows which features drive the fitted separation (Fig. 4**C2**). A loading is the weight that a component assigns to a feature: a larger absolute loading means a larger contribution to that component. In this example, the five features with the largest absolute loadings are the five signal features, V1–V5.

In the subject plot (Fig. 4**C1**), group A lies on the negative side of Component 1. V3 has the largest negative loading, which matches its higher mean trajectory in group A compared with group B; the same applies to V1 and V4. V2 and V5 have positive loadings, with higher mean trajectories in group B than in group A. This component is built on the temporal face that represents the average across time points (face 1), so the loading signs describe group differences on average over time. The accompanying curves are group mean trajectories, shown for description only to highlight differences across time. The signs themselves depend on the orientation of the component.

### 2.6 Benchmarks: simulated and real-data validation

We assessed the performance of our tensor methods in simulations with known temporal signals based on subject-level predictive validation, as well as in three longitudinal multi-omics cohorts (described in Table 2). The case studies in Section 2.7 separately illustrate biological interpretation.

**Table 2.** Real longitudinal cohorts used for predictive validation. Subjects are complete cases observed at every time in both blocks. Blocks give the omics type and the number of features before filtering; the numbers analysed after filtering are in Table 5.

| **Cohort** | **Outcome / design** | **Blocks** | **Subjects** | **Time points** | **Reference** |
|---|---|---|---|---|---|
| CMI-PB | Pertussis booster vaccination time course; aP vs wP priming | 31 antibody titres; 6,650 genes | 90 | 5 (days 0, 1, 3, 7, 14) | Shinde et al. (2024); da Silva Antunes et al. (2021) |
| HE INFANTS MFGM | Infant-formula RCT; diet arm (3 classes) | 295 genera; 120 metabolites | 52 | 4 (months 2, 4, 6, 12) | He et al. (2019) |
| MARS IBS | Observational IBS cohort; healthy, IBS-C, IBS-D | 2,895 genera; 43 metabolites | 46 | 6 (months 0–5) | Mars et al. (2020) |

#### 2.6.1 Simulation design

In all three simulations, we designed the groups with the same time-average, so that the group difference lay entirely in temporal shape. The generative models and parameter grids are detailed in Appendix B of S1 Appendix. In brief, feature *j* of subject *i* at time *t* was generated as *x*_*ijt*_ = *µ*_*j*_ + *a*_*j*_*z*_*g*(*i*)_(*t*) + *b*_*ij*_ + *e*_*ijt*_, where *z*_*g*_(*t*) is the temporal contrast of group *g* (with the same time-average in both groups), *a*_*j*_ = *±δ* for informative features and 0 otherwise (*δ* is the effect size), *b*_*ij*_ is a time-constant subject-by-feature offset (its standard deviation is the between-subject variation), and *e*_*ijt*_ is residual noise, independent or AR(1)-correlated. Each scenario was repeated on ten simulated datasets.

- **Simulation A: single-block temporal-shape recovery**. One assay contained linear or quadratic group trajectories. We assessed classification, recovery of the informative features and recovery of the temporal contrast.
- **Simulation B: representation, compression and basis scaling**. Three assays contained contrasts aligned with one discrete cosine transform (DCT) face or distributed across several faces. We compared our tensor methods against matrix-based methods with either concatenated time points or truncated DCT coefficients, with a separate diagnostic for column scaling.
- **Simulation C: multiblock temporal disagreement**. Omics 1 retained a fixed linear contrast while the contrasts in omics 2 and 3 were rotated from agreement to orthogonality. We assessed classification, feature recovery and agreement between the fitted and true temporal patterns.

The four feature classes and temporal contrasts are defined in Supplemental Table B.1 and Sections B.2–B.3 and illustrated in Supplemental Figs. B.1–B.3 of S1 Appendix.

#### 2.6.2 Real-data cohorts

We analysed three longitudinal multi-omics cohorts, each with two omic types (Table 2). Each block was arranged as subjects *×* features *×* time points (or visits), and each subject had one label across all time points. Only subjects observed at every time in both omic types were retained.

Briefly, taxa in HE INFANTS and MARS were centred log-ratio transformed, and their metabolite abundances were log_2_ transformed. CMI-PB gene-expression counts had been batch-corrected using ComBat-seq (Zhang et al., 2020) by the authors. Further details are provided in Supplemental Text C.

#### 2.6.3 Comparison with other methods

We chose methods according to their ability to perform unsupervised or supervised analyses and handle one or several omic types (Table 3). Matrix-based methods used one row per subject and one column per feature-time combination, i.e., time points were concatenated (PCA and PLS-DA for single-block comparisons, and block PLS-DA and DIABLO (Singh et al., 2019) for multiblock classification).

**Table 3.**
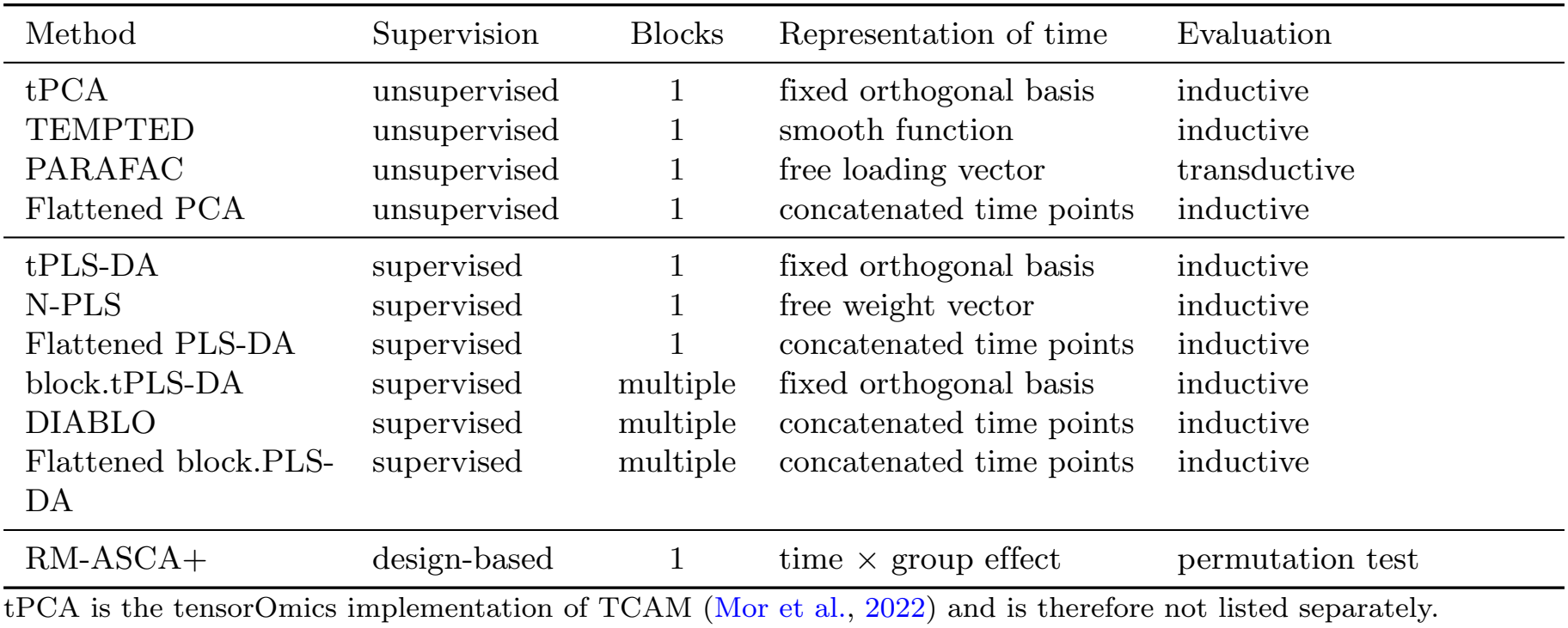
Comparator methods and evaluation protocols. In *inductive* evaluation, models are fitted using training subjects only, and held-out trajectories are scored using the fitted representation. In *transductive* evaluation, an unlabelled decomposition is fitted jointly to training and held-out measurements, without using held-out labels. Methods without a native classification rule use a nearest-centroid classifier trained on the training subjects’ scores and labels. Because held-out measurements contribute to a transductive representation, its error is not strictly out-of-sample and may be optimistic. RM-ASCA+ estimates population-level group-by-time effects and was evaluated using a permutation test.

The unsupervised temporal comparators differ in their representation of time: tPCA selects from a fixed orthogonal basis, TEMPTED (Shi et al., 2024) estimates a smooth function, and PARAFAC (Harshman et al., 1970) estimates a free temporal loading. TEMPTED was fitted with three components, a smoothing penalty of 10^*−*5^ and a 51-point time grid, and held-out subjects were scored with its own estimator, which re-estimates the subject loading with the feature and temporal loadings held fixed.

tPCA is the R implementation of TCAM (Mor et al., 2022) within tensorOmics.

MEFISTO (Velten et al., 2022) was not evaluated because the available implementation can interpolate unseen time points but cannot project an unseen subject.

For the supervised temporal comparators, N-PLS (Bro, 1996) estimates a free temporal weight rather than selecting a fixed basis element.

Finally, repeated-measures ANOVA simultaneous component analysis+ (RM-ASCA+) (Madssen et al., 2021) tests the group-by-time interaction using 199 label permutations. The method assesses a population-level effect rather than subject classification.

##### Scaling

Besides the omic-specific transformations applied in the real-data studies as pre-processing, we used centring but no further feature standardisation. Additionally, we performed a sensitivity analysis of feature scaling, in which each feature-time column was divided by its training-fold standard deviation. For the matrix-based comparators, we used their default feature scaling unless scaling was explicitly disabled for the Simulation B diagnostic.

##### Prespecified settings

Three components were fixed *a priori* for our tensor methods and the component-based comparators alike and were not tuned within cross-validation. The tensor methods used the complete DCT basis of the visits. DIABLO was configured to select 30 columns per omic block and component.

For feature recovery, we considered the features with the largest absolute loading value per component for each method. We then examined the top 15 of the 200 features in Simulation A, which has 15 informative features, and the top 30 in each assay in Simulations B and C, where each assay includes 30 discriminative features. Selection stability was measured as the overlap of the selected feature sets across folds. In Simulation B the matrix-based comparators retained either 2, 3 or all 6 DCT coefficients. In Simulation C we compared block-to-block design weights of 0 and 0.2, and in the real-cohort multiblock analyses weights of 0, 0.5 and 1. The complete simulation grid is in Supplemental Table B.2.

#### 2.6.4 Evaluation measures

Classification performance was measured by balanced error rate (BER), the mean of the per-class error rates, to account for unbalanced subjects per class (Rohart et al., 2017). Throughout the simulations and real cohorts, validation was performed at subject level: all time points from each held-out subject were excluded from training. Five-fold cross-validation was repeated ten times, with the same subject splits used for all methods within a dataset or cohort.

Table 3 summarises how scores were obtained for held-out subjects. The *inductive* and *transductive* protocols differ in whether held-out measurements contribute to fitting the representation. Methods without a native classification rule, including all unsupervised methods, were evaluated using the same nearest-centroid classifier. This classifier was fitted after decomposition using only the training subjects’ scores and labels. Thus, differences in classification performance primarily reflect the fitted representations, while the unsupervised decompositions remain label-free.

Under the *inductive* protocol, the model was fitted using only the training subjects and, for supervised methods, their labels. Held-out subjects were then projected using the fitted feature and temporal loadings.

TEMPTED additionally estimated a subject-specific loading for each held-out subject while keeping the feature and temporal loadings fixed; the other methods used their native projection or prediction functions.

The resulting error estimates predictive performance in new subjects.

Under the *transductive* protocol, an unsupervised representation was fitted jointly to the measurements from training and held-out subjects, without using held-out labels. Within each fold, the classifier was still trained using only the training subjects’ scores and labels. This protocol was required for PARAFAC because the implementation used could not project new subjects onto a decomposition fitted using training subjects alone. It applies only to unsupervised methods, as fitting a supervised model jointly to all subjects would introduce overfitting. Because held-out measurements contribute to the fitted loadings, the resulting error is not strictly out-of-sample and may be optimistic relative to an inductive estimate.

To quantify the effect of the evaluation protocol, we evaluated unsupervised methods that support both protocols using identical folds.

### 2.7 Case studies

Three case studies illustrate the biological application and interpretation of tensorOmics across different study designs and omic blocks. Table 4 summarises their sample sizes, time points and post-filtering feature counts.

**Table 4.**
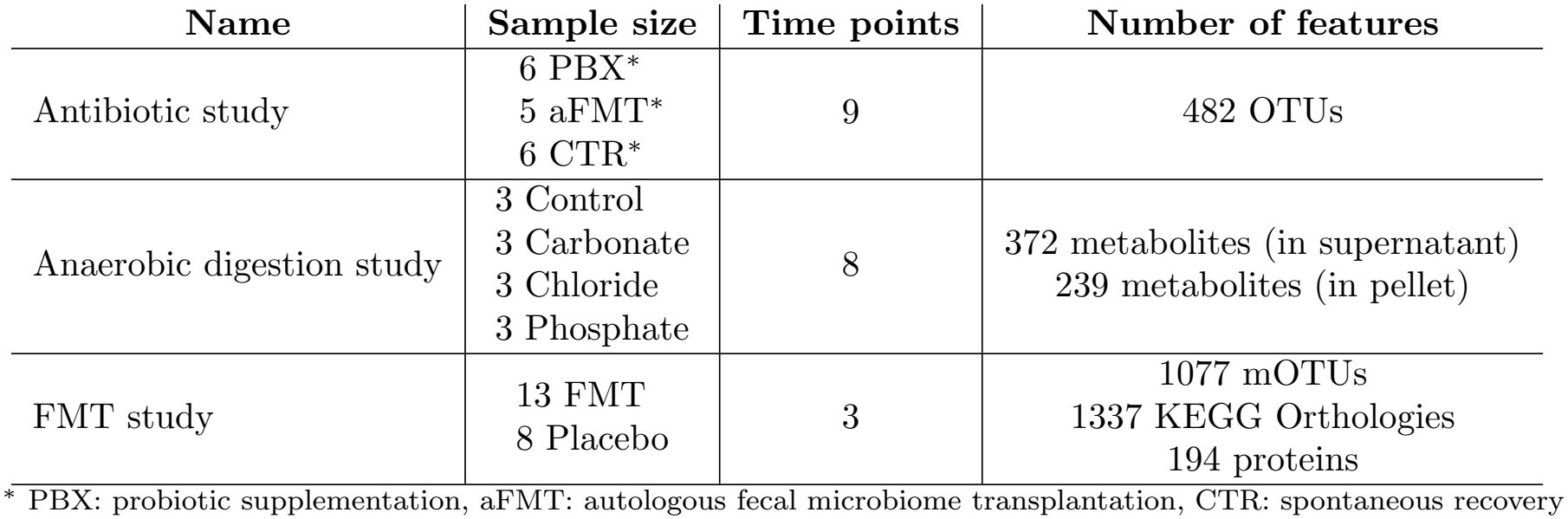
Case study designs. Sample sizes, time points and feature counts after filtering.

| Name | Sample size | Time points | Number of features |
|---|---|---|---|
| Antibiotic study | 6 PBX<sup>*</sup><br>5 aFMT<sup>*</sup><br>6 CTR<sup>*</sup> | 9 | 482 OTUs |
| Anaerobic digestion study | 3 Control<br>3 Carbonate<br>3 Chloride<br>3 Phosphate | 8 | 372 metabolites (in supernatant)<br>239 metabolites (in pellet) |
| FMT study | 13 FMT<br>8 Placebo | 3 | 1077 mOTUs<br>1337 KEGG Orthologies<br>194 proteins |

<sup>\*</sup> PBX: probiotic supplementation, aFMT: autologous fecal microbiome transplantation, CTR: spontaneous recovery

#### 2.7.1 Case study 1: Antibiotic study (single-omic, classification)

Suez et al. (2018) studied gut microbiome reconstitution after antibiotics in healthy subjects assigned to 21-day probiotic supplementation (PBX), autologous fecal microbiome transplantation (aFMT; FMT in the Results and figures) using pre-antibiotic samples, or spontaneous recovery (CTR). Stool samples were collected at baseline (BAS; days 3 and 6), during antibiotics (ABX; days 10 and 13), and during intervention (INT; days 17, 20, 28, 35 and 42).

The original study removed operational taxonomic units (OTUs) with relative abundances below 0.1% and rarefied samples to 10,000 reads. We used the dataset provided in TCAM, which additionally applied log-fold baseline normalisation (Mor et al., 2022).

We fitted tPLS-DA using the complete DCT-II basis and three components. The subject-level class response was supplied unchanged to every face, deflated globally after each component, and faces were compared after Frobenius scaling. No additional feature standardisation was applied. The displays report the first two components and the ten largest absolute feature loadings for each.

#### 2.7.2 Case study 2: Ammonia study (two-omic, unsupervised)

Wang et al. (2024) examined the metabolic effects of ammonia sources on anaerobic digestion in 12 digesters: three uninhibited controls and three receiving each of 14.9 g ammonium carbonate, 16.6 g ammonium chloride, or 20.5 g ammonium hydrogen phosphate. Liquid samples collected on days 0, 6, 13, 20, 27, 34, 41, 48 and 55 were separated into supernatant and pellet fractions.

Metabolites contributing less than 0.01% of total intensity were removed, and the remaining features were log-transformed after adding one. Because of high variability between days 0 and 6, we analysed day 6 onwards.

We fitted tPLS with canonical deflation to the supernatant and pellet tensors, without a response, using the complete DCT-II basis and four components. Four components were retained because the cross-block covariance carried by each component, measured by its singular value, decreased sharply up to the fourth component and only gradually afterwards. The first three components were built on the average over time, and the fourth was the first to use a temporal contrast. This criterion is descriptive and was not cross-validated. No additional feature standardisation or block scaling was applied. The displays report the first two components and the eight largest absolute loadings per fraction; correlations were calculated for all 64 cross-fraction pairs among these features, and pairs with mean absolute correlation above 0.6 were displayed. The first two components did not depend on the number of components fitted.

#### 2.7.3 Case study 3: FMT study (multi-omic, classification)

Díaz-García et al. (2024) compared repeated low-dose fecal microbiota transplantation (FMT) with placebo in patients with HIV. At baseline and weeks 1, 8 and 24, they measured 345 inflammatory plasma proteins by proximity extension assay as normalised protein expression values on a log_2_ scale. Shotgun metagenomics profiled species-level metagenomic operational taxonomic units (mOTUs) and functional Kyoto Encyclopedia of Genes and Genomes orthology (KO) terms.

We analysed baseline and weeks 8 and 24 because placebo microbiome data were unavailable at week 1. Week 8 marked the end of dosing, and week 24 was 16 weeks later. Features contributing less than 0.01% of total abundance were removed. After adding one, mOTUs were centred log-ratio transformed and KOs were log-transformed; proteins were scaled as described by Díaz-García et al. (2024). Systematic variation in the mOTU and KO blocks was corrected as detailed in Supplemental Section E (Fig. E.1; Table E.1). No feature standardisation was applied to any of the three blocks.

The three layers formed separate arrays: 21*×*1,077*×*3 for mOTUs, 21*×*1,337*×*3 for KOs and 21*×*194*×*3 for proteins. We applied block.tPLS-DA to integrate the tensors by their shared subjects, time and treatment groups, with three components. In our reported analysis, we assigned a design weight of 1 to all omic–omic and omic–outcome pairs. We provide an additional sensitivity analysis (Supplemental Section E) for an omic–omic weight of 0.1.

## 3 Results

### 3.1 Benchmark results

We first report the results from the three simulation schemes, followed by the classification of held-out subjects in three longitudinal multi-omics cohorts. Additional results are reported in Appendix D of S1 Appendix.

#### 3.1.1 Simulation A: single-block temporal-shape recovery

The two groups in Simulation A had the same time-average for every feature and differed only in trajectory shape. As the effect size increased, the BER of tPLS-DA decreased from 0.450 at effect size *δ* = 0.05 to 0.068 at *δ* = 0.15 and 0.000 at *δ* = 0.3 for the linear shape. We observed a similar pattern for the quadratic shape (Fig. 5**A1–A2**). PLS-DA on flattened visits had a slower decrease, whereas N-PLS, TEMPTED and PARAFAC remained near chance (Supplemental Table D.1).

**Fig 5.**
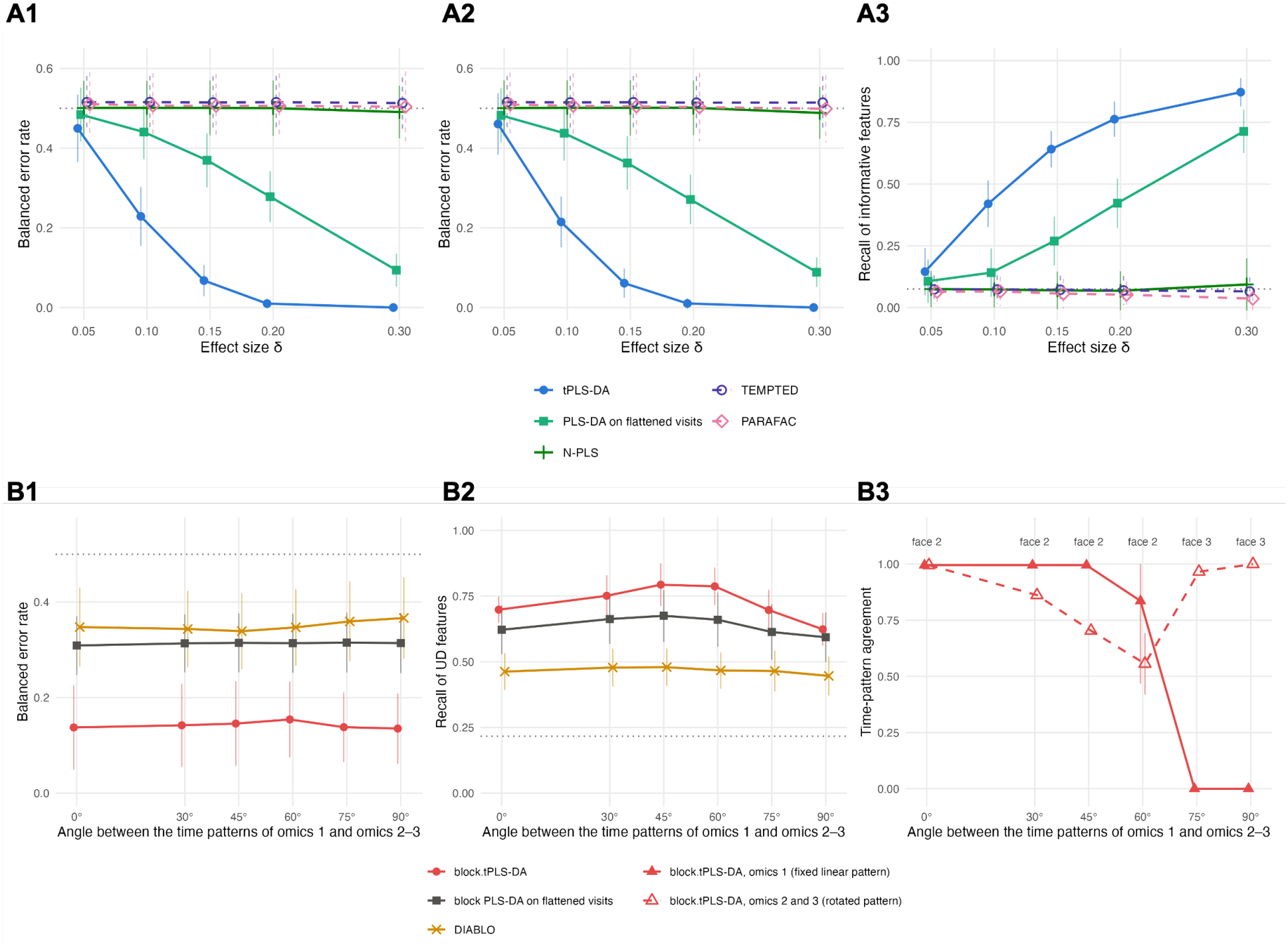
Benchmark of tensor partial least squares discriminant analysis (tPLS-DA) and its multiblock extension (block.tPLS-DA) for group differences in temporal shape. **A1–A3**: Simulation A (single-omic). The groups have the same time-average and differ only in trajectory shape; *δ* indicates effect size. **A1–A2**: balanced error rate (BER) for linear and quadratic contrasts. **A3**: recall of the 15 informative features among the 15 top-ranked for the linear contrast. **B1–B3**: Simulation C (multi-omics). Omics 1 has a fixed linear contrast, while the contrast in omics 2 and 3 is rotated from shared (0°) to orthogonal (90°). **B1**: BER at *δ* = 0.1. **B2**: recall at *δ* = 0.3 of uncorrelated discriminative (UD) features among the 30 top-ranked per omic. **B3**: absolute correlation at *δ* = 0.3 between the fitted temporal pattern and the true patterns in omics 1 and in omics 2–3; labels indicate the face selected most often (face 2, steady trend; face 3, U-shaped). Points are means across 10 simulated datasets, bars show one standard deviation and dotted lines indicate chance.

Face scaling was important when stable between-subject variation was present. At *δ* = 0.2, scaled tPLS-DA retained a BER of 0.010 across between-subject standard deviations from 0 to 2. Without face scaling, BER reached chance (0.502) at a standard deviation of 1 or more (Supplemental Table D.2). The error also remained low across the tested cohort sizes and residual correlations.

tPLS-DA recovered both the informative features and their temporal pattern. At *δ* = 0.2, 0.763 of its top 15 features were informative, compared with 0.423 for PLS-DA on flattened visits and 0.068 for N-PLS (chance 0.075; Fig. 5**A3**). It also selected fewer null features (false-positive rate 0.019, against the same chance level of 0.075) and its selections were the most reproducible across folds (Jaccard overlap 0.480, against 0.039 for random sets of 15 features), as detailed in Supplemental Table D.3.

#### 3.1.2 Simulation B: representation of time and basis scaling

Our Simulation B identified two limitations in interpreting temporal representations.

First, each tensor component used one time face. block.tPLS-DA recovered the linear and quadratic contrasts, both of which lay on one face (agreement 0.995 and 0.982). For off-basis-smooth and transient contrasts distributed across several faces, it reached the maximum or near-maximum agreement available from one face (0.634 and 0.721), while retaining low BER (0.084 and 0.076 at *δ* = 0.1; Supplemental Table D.4). The fitted time pattern was therefore interpreted as the dominant temporal direction, not necessarily the complete contrast.

Second, pre-processing alone can create an apparent performance difference between otherwise equivalent representations. The full discrete cosine transform of the six visits is an exact orthogonal rotation of the flattened visit representation and therefore preserves the same information. With the default column scaling in block PLS-DA (matrix-based), the flattened and transformed representations yielded classification errors of 0.297 and 0.102, respectively. When column scaling was disabled, both yielded an error of 0.310 (Supplemental Table D.5). The observed difference therefore arose from the interaction between representation and scaling. We consequently interpreted Simulations B and C as analyses of method behaviour and do not use them to rank tensor and matrix methods unless their scaling was matched.

#### 3.1.3 Simulation C: multiblock temporal disagreement

In Simulation C, omics 2 and 3 encoded the group difference with a time pattern rotated away from the fixed linear pattern of omic 1. The BER of block.tPLS-DA remained stable as disagreement increased: at *δ* = 0.1 and design weight 0, it was 0.138 at 0° and 0.135 at 90° (Fig. 5**B1**; Supplemental Table D.6). Increasing the omic–omic design weight from 0 to 0.2 gave a higher BER than design weight 0 (Supplemental Table D.6), so linking the omics did not improve classification when they disagreed about the temporal pattern. The matrix-based methods were also stable across angles using column scaling (their errors are not directly comparable against block.tPLS-DA, as highlighted above). At *δ* = 0.3, block.tPLS-DA recovered 0.698 of the UD features at 0° and 0.624 at 90° (chance 0.217; Fig. 5**B2**; Supplemental Table D.7).

Classification remained accurate even when the temporal interpretation changed. Up to 45°, the first component followed the pattern of omics 1; at 75° and 90°, it followed the shared pattern of omics 2 and 3 instead, and agreement with omics 1 reached zero (Fig. 5**B3**). The classification error nevertheless remained at most 0.012 at *δ* = 0.3 (Supplemental Table D.6). Because omics 2 and 3 were rotated together, this switch reflected a two-against-one construction rather than a general threshold. Thus, strong classification could conceal the loss of an omics-specific time pattern.

#### 3.1.4 Longitudinal cohorts

##### Single omics

We found that performance varied between supervised methods depending on the cohort and the omics type (Table 5). When features were scaled, a tensor method (tPLS-DA or N-PLS) was best in six out of twelve single omics datasets and PLS-DA on flattened time in six; several differences were small relative to variation across cross-validation repeats. Scaling changed the best method in three of the six blocks, but no method was best throughout (Supplemental Table D.8). Feature overlap across folds exceeded its random-set expectation, but no method was consistently the most stable; among the three supervised methods, tPLS-DA was the least stable in three of four unscaled blocks and all four scaled blocks assessed (Supplemental Table D.9). Unsupervised methods were reported as a separate family and were not ranked against the supervised methods.

**Table 5.** Longitudinal cohorts: single-block classification. Balanced error rate (BER), mean (SD) over 10 repeats of 5-fold subject-level cross-validation, with three components and unscaled data. Numbers of features are indicated in brackets. The lowest error among the supervised rows, and among the unsupervised rows in each column, is shown in bold. *PLS-DA on flattened visits* does not model the order of visits. Unsupervised methods were treated as a separate family and were not ranked against the supervised rows. Lower values indicate better classification; chance was 0.500 for CMI-PB (two classes) and 0.667 for HE INFANTS and MARS (three classes).

|  | CMI-PB (*n* = 90) |  | HE INFANTS (*n* = 52) |  | MARS (*n* = 46) |  |
|---|---|---|---|---|---|---|
| Method | titres (31) | genes (1000) | genera (124) | metabolome (101) | genera (301) | metabolome (26) |
| *Supervised* |  |  |  |  |  |  |
| tPLS-DA | **0.309 (0.029)** | 0.395 (0.042) | 0.600 (0.054) | 0.407 (0.070) | 0.559 (0.047) | **0.409 (0.082)** |
| N-PLS | 0.321 (0.041) | 0.372 (0.027) | **0.489 (0.050)** | 0.387 (0.029) | 0.649 (0.052) | 0.546 (0.078) |
| PLS-DA on flattened visits | 0.318 (0.031) | **0.348 (0.039)** | 0.575 (0.048) | **0.366 (0.037)** | **0.558 (0.031)** | 0.506 (0.049) |
| *Unsupervised (separate family)* |  |  |  |  |  |  |
| tPCA | **0.277 (0.012)** | 0.368 (0.023) | **0.510 (0.071)** | 0.315 (0.017) | **0.500 (0.070)** | **0.441 (0.031)** |
| PCA on flattened visits | 0.278 (0.018) | 0.377 (0.022) | 0.511 (0.056) | **0.284 (0.019)** | 0.512 (0.053) | 0.462 (0.025) |
| TEMPTED | **0.277 (0.014)** | **0.362 (0.022)** | 0.592 (0.056) | 0.297 (0.033) | 0.517 (0.037) | 0.514 (0.044) |

##### Multiblock

We found that block.tPLS-DA had the lowest BER across design weights for CMI-PB and MARS, whereas no method clearly dominated on HE INFANTS (Table 6). Combining blocks improved on the best supervised single-block result for CMI-PB (0.247 versus 0.309) and HE INFANTS (0.350 versus 0.366), but not for MARS (0.433 versus 0.409). These comparisons were not matched for scaling, as block.tPLS-DA centred but did not scale the data, whereas block PLS-DA and DIABLO used default feature scaling.

**Table 6.** Longitudinal cohorts: multiblock classification. Balanced error rate (BER), mean (SD) over 10 repeats of 5-fold subject-level cross-validation, two omics blocks per cohort, with three components and unscaled data. The design weight links the omics: a weight of 0 only links each block to the outcome, whereas a weight of 1 links every pair of blocks. The lowest error in each row is shown in bold. block.tPLS-DA centred but did not scale the data and assigned a new subject to the nearest class centroid; block PLS-DA and DIABLO were run with default column scaling and the mixOmics package’s own classification rule. Lower values indicated better classification; chance was 0.500 for CMI-PB and 0.667 for HE INFANTS and MARS.

| Cohort | Design weight | block.tPLS-DA | block PLS-DA, flattened time | DIABLO |
|---|---|---|---|---|
| CMI-PB | 0 | **0.255 (0.017)** | 0.313 (0.024) | 0.299 (0.050) |
|  | 0.5 | **0.247 (0.024)** | 0.361 (0.035) | 0.330 (0.044) |
|  | 1 | **0.251 (0.020)** | 0.376 (0.029) | 0.355 (0.039) |
| HE INFANTS | 0 | 0.378 (0.038) | 0.372 (0.023) | **0.371 (0.037)** |
|  | 0.5 | 0.401 (0.036) | **0.384 (0.028)** | 0.386 (0.048) |
|  | 1 | 0.387 (0.030) | **0.350 (0.026)** | 0.388 (0.034) |
| MARS | 0 | **0.433 (0.037)** | 0.531 (0.036) | 0.594 (0.044) |
|  | 0.5 | **0.484 (0.023)** | 0.545 (0.053) | 0.559 (0.034) |
|  | 1 | **0.487 (0.030)** | 0.526 (0.050) | 0.520 (0.037) |

Overall, in these large longitudinal multi-omics studies, our tensor methods classified held-out subjects about as well as the established methods, with no method performing best in every setting. However, the methods highlight temporal patterns at the feature level, as illustrated in the case studies below.

### 3.2 Case study results

#### 3.2.1 Case study 1: Supervised tensor analysis identified treatment-specific microbial signatures in longitudinal microbiome data

tPLS-DA separated the antibiotic-treated participants assigned to autologous fecal microbiota transplantation (FMT), probiotics (PBX), or spontaneous recovery (CTR) in the fitted subject scores (Fig. 6**A–B**). Component 1 separated PBX from CTR and FMT. PBX participants generally had more negative scores on this component than CTR and FMT participants (Fig. 6**B**). *Erysipelatoclostridium ramosum* was the only top-ranked taxon with a higher trajectory in PBX; the other nine taxa recovered more strongly in CTR and FMT and included several bacteria previously associated with microbiome recovery, such as *Bacteroides ovatus* and *Roseburia inulinivorans* (Fig. 6**C1**) (Chen et al., 2023). *Bacteroides thetaiotaomicron*, also among the top-ranked taxa for Component 1, was one of the recovery-associated bacteria that promoted recovery in a mouse model in a cross-cohort metagenome-wide association study (Chng et al., 2020). The weaker recovery of these commensals under PBX is consistent with a previous human study that compared the same three post-antibiotic strategies and reported delayed reconstitution of the indigenous gut microbiome with probiotics relative to spontaneous recovery and rapid reconstitution with autologous FMT (Suez et al., 2018).

**Fig 6.**
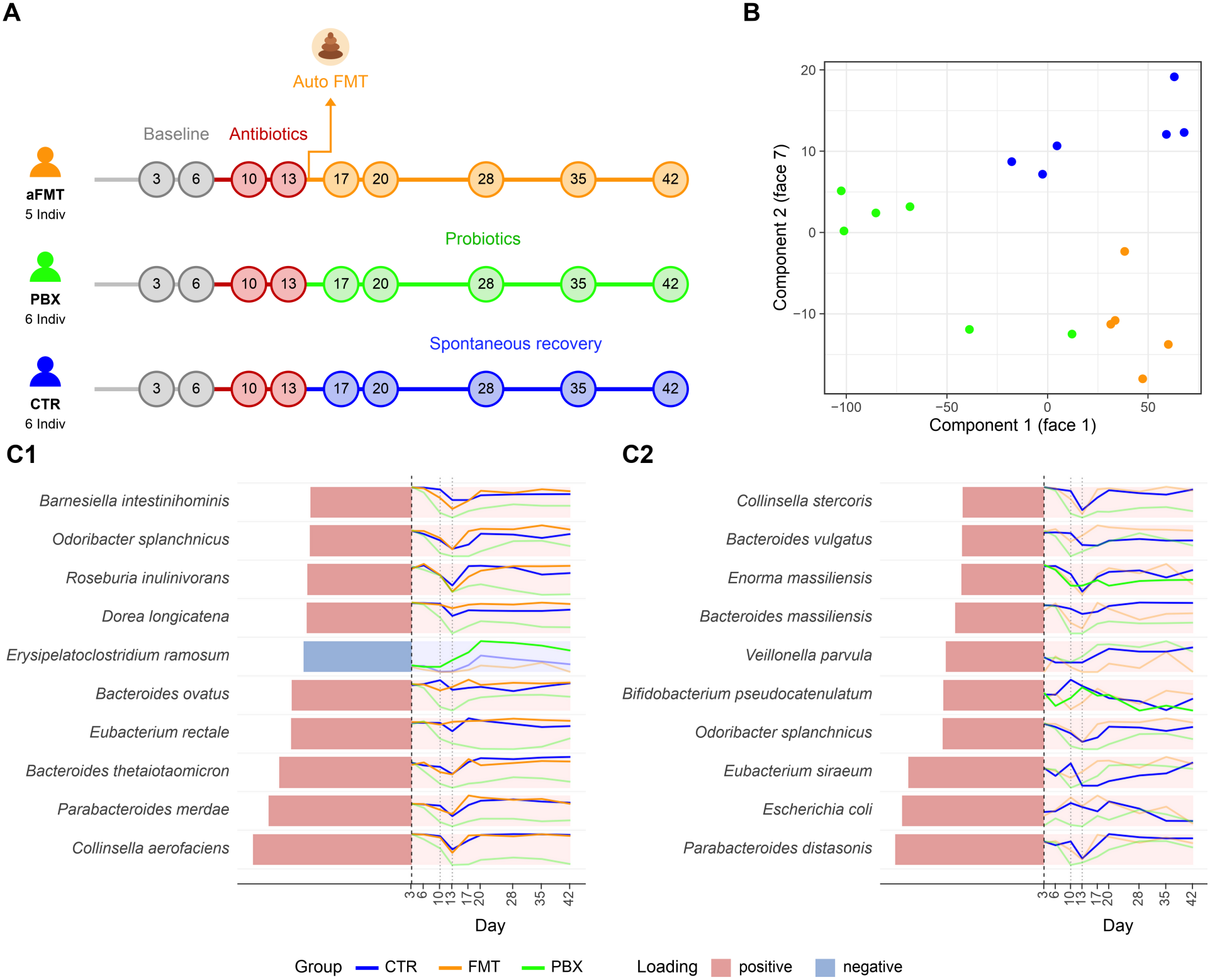
Tensor partial least squares discriminant analysis (tPLS-DA) of microbiome recovery after antibiotics in humans. **A**: study design and sampling schedule for participants assigned after antibiotics to autologous fecal microbiota transplantation (aFMT; FMT in the figure, *n* = 5), probiotics (PBX, *n* = 6), or spontaneous recovery (CTR, *n* = 6). **B**: fitted subject scores on Components 1 and 2; each point represents one participant’s complete trajectory. **C1–C2**: the ten taxa with the largest absolute loadings and their group-specific mean trajectories for Components 1 and 2, respectively. Because group labels are used to fit tPLS-DA, separation in the fitted scores is descriptive.

Component 2 highlighted a fast-changing temporal term, separating CTR from FMT, with PBX in intermediate positions (Fig. 6**B, C2**). All ten top-ranked taxa for this component had positive loadings (Fig. 6**C2**). These included *Escherichia coli*, a member of the *Enterobacteriaceae*, a family that often expands rapidly in relative abundance soon after broad-spectrum antibiotics in healthy adults (Palleja et al., 2018), which is compatible with a fast-changing temporal term. Because group labels are used to fit tPLS-DA, the separation in the fitted scores remains descriptive. With 17 subjects, leave-one-subject-out refits retained the selected time term in only 8 of 17 refits for Component 1 and 10 of 17 for Component 2, and recovered fewer than five of the ten top taxa on average. However, low stability is not unexpected because post-antibiotic recovery is individualised and can remain incomplete in some subjects (Dethlefsen and Relman, 2011, Palleja et al., 2018). The named taxa are therefore best treated as hypotheses for follow-up in larger cohorts rather than as validated treatment-specific markers. With only 17 subjects, Case Study 1 illustrates how the fitted components and loadings are interpreted and does not assess predictive performance, which we evaluate in held-out subjects in the benchmarks (Section 3.1).

#### 3.2.2 Case study 2: Unsupervised tensor integration suggested metabolic coordination across supernatant and pellet fractions

tPLS linked the longitudinal metabolomes of the supernatant and pellet fractions. Both blocks showed similar treatment structure: Component 1 separated Control and Chloride from Carbonate and Phosphate, while Component 2 provided additional discrimination (Fig. 7**B1, C1**). The pellet showed greater overlap between Control and Chloride than the supernatant.

**Fig 7.**
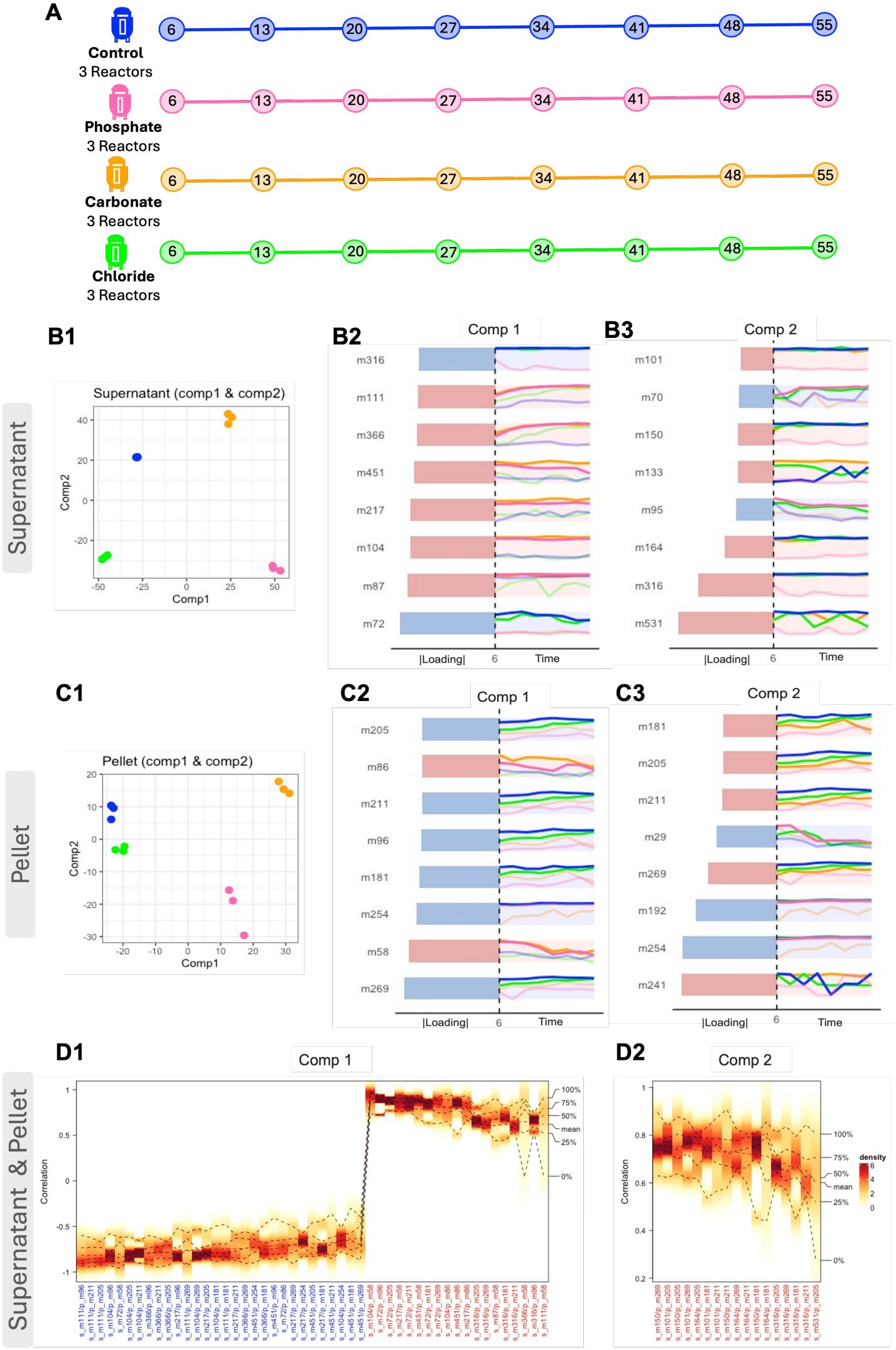
Tensor partial least squares (tPLS) integration of supernatant and pellet metabolomes under ammonium treatments in bioreactors. **A**: study design and weekly sampling of 12 anaerobic digesters assigned to control, ammonium carbonate, ammonium chloride or ammonium hydrogen phosphate conditions. **B1–B3**: fitted subject scores (B1) and the eight metabolites with the largest absolute loadings for Components 1 (B2) and 2 (B3) in the supernatant. **C1–C3**: corresponding fitted scores (C1) and loading plots (C2–C3) for the pellet. **D1–D2**: temporal correlations between supernatant and pellet metabolite pairs formed from the top-eight sets for Components 1 and 2, respectively; pairs with mean absolute correlation greater than 0.6 were retained. Prefixes `s_` and `p_` denote supernatant and pellet metabolites.

The loading profiles linked the treatment separation in the subject plots to distinct metabolite patterns in each compartment. On Component 1, six of the eight leading supernatant metabolites (m87, m104, m217, m451, m366 and m111) had positive loadings and aligned with the Carbonate and Phosphate groups, whereas m72 and m316 had negative loadings and aligned with Control and Chloride (Fig. 7**B2**). Their trajectories included both stable group differences, as observed for m104 and m316, and treatment differences that changed over time, as observed for m111 and m366. Component 2 captured weaker, more treatment-specific patterns: m316, m164, m150 and m101 were lower in Phosphate, whereas m95 and m70 were higher in Phosphate and Chloride (Fig. 7**B3**); the pattern for m95 was consistent with Wang et al. (2024).

The pellet loadings showed complementary patterns. Six of the eight leading contributors to Component 1 (m269, m254, m181, m96, m211 and m205) had negative loadings and aligned with Control and Chloride, whereas m58 and m86 had positive loadings and declined over time in Carbonate and Phosphate (Fig. 7**C2**). Component 2 again showed weaker treatment effects: m254 and m192 remained relatively abundant in all groups except Carbonate, while metabolites with positive loadings tended to have their lowest abundance in Phosphate (Fig. 7**C3**). Thus, the two components captured a dominant contrast between treatment groups together with weaker, treatment-specific temporal variation across the supernatant and pellet compartments.

To examine the corresponding cross-compartment associations, we paired the eight metabolites with the largest absolute loadings from each compartment, giving up to 64 candidate pairs per component, and retained pairs with an average absolute correlation above 0.6. Component 1 pairs showed predominantly strong and temporally stable positive correlations, compatible with persistent coordination such as parallel production or coupled consumption across compartments (Fig. 7**D1**). Negative correlations were less stable and more dispersed. Component 2 correlations were weaker and more variable over time, consistent with the smaller amount of shared variation represented by this component (Fig. 7**D2**).

#### 3.2.3 Case study 3: Multiblock integration of taxonomic, functional and protein profiles following FMT

We applied block.tPLS-DA to integrate the taxonomic (mOTU), functional (KO) and plasma protein profiles through two temporal patterns. Component 1 represented the average over the three time points, whereas Component 2 contrasted week 8 with baseline and week 24 (Fig. 8**B1–B3**).

**Fig 8.**
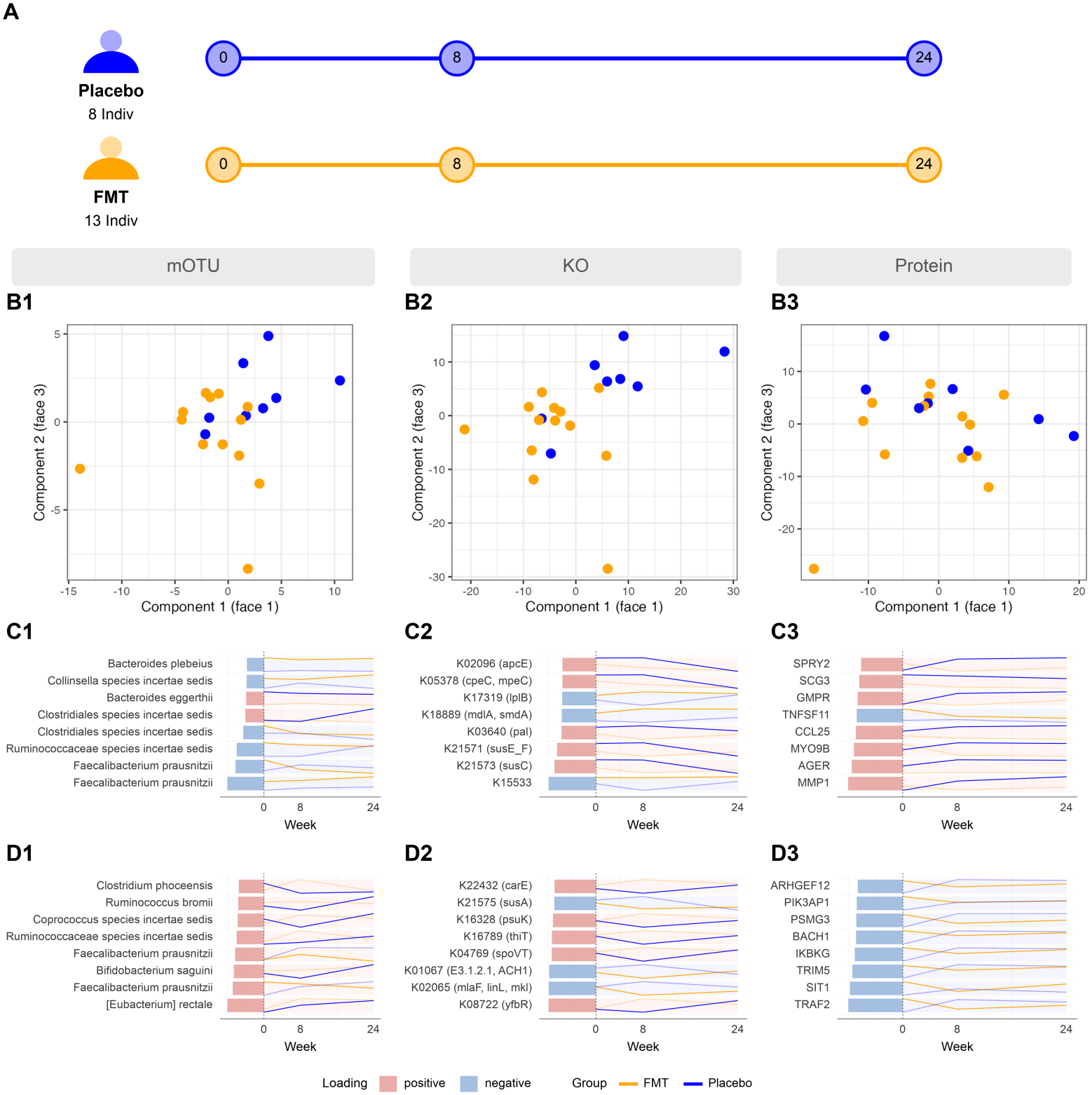
Multiblock tensor partial least squares discriminant analysis (block.tPLS-DA) of taxonomic, functional and protein profiles in fecal microbiota transplantation (FMT) and placebo recipients. **A**: study design and sampling of 13 FMT and 8 placebo recipients at baseline, the end of dosing (week 8) and follow-up (week 24). The blocks comprised metagenomic operational taxonomic units (mOTUs), Kyoto Encyclopedia of Genes and Genomes orthology terms (KOs) and plasma proteins; the mOTU and KO blocks were batch-corrected. **B1–B3**: fitted subject scores for the mOTU, KO and protein blocks. Component 1 uses the average over time (face 1), and Component 2 contrasts week 8 with the other time points (face 3). **C1–C3** and **D1–D3**: the eight features with the largest absolute loadings and their group-specific mean trajectories for Components 1 and 2, respectively. Repeated taxon names denote distinct mOTUs. Because treatment labels are used to fit the components, separation in B1–B3 is expected; held-out classification and stability are reported in the text.

On Component 1, FMT recipients had higher average abundance of two *Faecalibacterium prausnitzii* mOTUs and a Ruminococcaceae species (Fig. 8**C1**), consistent with donor selection and reported engraftment in the trial (Díaz-García et al., 2024, Serrano-Villar et al., 2021). KO loadings contrasted functions related to complex carbohydrate metabolism and nutrient transport in FMT recipients with starch-utilisation functions in placebo recipients (Fig. 8**C2**). Seven of the eight largest protein loadings, including CCL25, AGER and MMP1, were higher on average in placebo recipients (Fig. 8**C3**). However, Component 1 also captured pre-treatment differences, including one *F. prausnitzii* mOTU that was already more abundant in FMT recipients at baseline.

Component 2 described a transient week-8 microbial pattern. Several KO functions, including the sporulation regulator SpoVT, increased at week 8 in FMT recipients and decreased in placebo recipients before returning towards baseline (Fig. 8**D2**). Its leading mOTUs included Lachnospiraceae and Ruminococcaceae taxa associated with the trial’s transient engraftment, although some contrasts reflected decreases in placebo recipients as much as increases after FMT (Fig. 8**D1**) (Serrano-Villar et al., 2021). A model-independent comparison supported the functional result: KO profiles differed between groups at week 8 (*p* = 0.004; Bonferroni-adjusted *p* = 0.036 across nine block-by-time comparisons), but not at baseline or week 24. Evidence was weaker for mOTUs (*p* = 0.09) and absent for proteins; we therefore did not interpret the Component 2 protein loadings, which were dominated by baseline differences (Fig. 8**D3**).

In repeated three-fold subject-level cross-validation, the three-block model had a BER of 0.266 (SD 0.065), compared with 0.521 under permuted labels (*p* = 0.03); the KO scores had the lowest block-specific error (0.263). Without batch correction, the BER was much higher at 0.465 (SD 0.098, see Supplemental Table E.2).

## 4 Discussion

Longitudinal multi-omics studies require methods that preserve repeated measurements while integrating information across molecular layers. Most existing integration methods were developed for cross-sectional data and require longitudinal profiles to be flattened, either by treating repeated visits as independent observations or by representing each visit as a separate set of features. Our tensorOmics framework instead retains each subject’s complete trajectory as the unit of analysis, represents time using a fixed orthogonal basis and constructs components that link temporal variation to molecular features. Within a common framework, it supports exploratory analysis, regression, classification and multiblock integration, including projection of new subjects. This framework extends predominantly unsupervised approaches, including the tensor methods TEMPTED (Shi et al., 2024) and PARAFAC (Harshman et al., 1970) and the multi-view method MEFISTO (Velten et al., 2022), as well as supervised approaches restricted to a single longitudinal block, such as N-PLS (Bro, 1996). It also complements methods designed to estimate population-level longitudinal effects, such as RM-ASCA+ (Madssen et al., 2021).

The simulations showed that this representation is particularly useful when biological groups differ in the shape of their trajectories rather than in their average levels. In these settings, our tensors recovered the relevant temporal contrasts and prioritised the contributing features. They also highlighted that apparent differences between tensor and matrix representations can arise from pre-processing rather than from the underlying information available to either method. Standardisation choices should therefore be specified, applied consistently across methods and estimated using only the training subjects within each resampling split.

Across the public cohorts, our tensor methods achieved held-out classification performance comparable with established approaches, although no method performed best in every setting. Thus, our methods retained longitudinal trajectories and provided interpretable temporal and feature loadings while maintaining comparable predictive performance. Multiblock integration improved performance in some cohorts but not others. Moreover, predictive performance alone did not establish whether a multiblock component represented a temporal pattern shared across all omics blocks. The selected temporal term and corresponding feature trajectories should therefore be examined within each block alongside held-out performance.

The case studies illustrate how the framework can generate biologically interpretable temporal patterns and candidate molecular signatures across diverse longitudinal designs. They also emphasise the distinction between fitted structure and independently supported biological evidence. Small sample sizes, instability of selected features, residual technical effects and correlations induced by the modelling objective can all limit interpretation. Accordingly, fitted components should be treated as a basis for generating hypotheses, with candidate findings evaluated using complementary evidence such as model-independent tests, resampling-based stability analyses, negative controls and external biological knowledge.

Several limitations affect the generalisability of the present findings. Each component currently captures a single dominant temporal pattern and may not represent all longitudinal variation. The implementation also requires a complete, aligned visit grid and does not directly accommodate unequal observation intervals. Interpolation may broaden applicability but introduces uncertainty, particularly when missingness is informative or intervals are long (Straube et al., 2015). In multiblock analyses, covariance-based objectives may give greater influence to blocks with higher post-transformation variance, and different omics technologies may require different preprocessing and scaling. These choices should therefore be stated explicitly. Finally, the benchmark covered a limited number of cohorts and comparators, and the limited stability of feature selection warrants caution when interpreting individual molecular features.

Tucker decomposition was not included as a comparator because its application would require additional choices regarding the number of components in each mode, model constraints and temporal regularisation (Tucker, 1966, Kolda and Bader, 2009). Moreover, multiblock datasets with different numbers of features cannot be represented as a single tensor. They would require a coupled Tucker formulation with shared subject and time modes but block-specific feature factors and core tensors (Acar et al., 2014, 2015). Developing and validating such a model was beyond the scope of this study. The PARAFAC result in Simulation A should therefore not be interpreted as representative of Tucker decompositions.

Future developments could allow components to combine multiple temporal terms or select block-specific terms, thereby accommodating responses distributed across several temporal patterns or occurring at different times across omics layers. Extensions to incomplete or partially overlapping cohorts, irregularly sampled data, nonlinear relationships and mixed-effects formulations would further broaden the framework. Larger longitudinal studies with more than two omics blocks will also be important for establishing when temporal multiblock modelling improves prediction and biological interpretation.

In summary, tensorOmics provides a unified framework for analysing longitudinal omics data at the level of the subject trajectory. It links temporal variation to contributing molecular features while supporting exploratory, predictive and multiblock analyses. Its principal value lies in combining these capabilities within an interpretable longitudinal representation; reliable biological conclusions nevertheless depend on careful pre-processing, held-out evaluation and independent assessment of the fitted temporal patterns and selected features.

## Declarations

### Code availability

tensorOmics is available as an R package at https://github.com/brendanlu/tensorOmics. Code used for the case studies in this manuscript can be accessed at https://github.com/lecaolab/lecaolab-paper-tensorOmics.

## Supporting information

**S1 Appendix**. Supplementary simulation models, evaluation protocols, benchmark results and batch-correction analyses.

