## Supplemental material for "tensorOmics: Data integration for longitudinal omics data using tensor factorisation"

### S1 Appendix

### Appendix A Detailed tPLS component extraction and deflation

Algorithm 2 gives the full two-block procedure underlying tPLS. At component $h$, $\mathbf{Z}_{h,k}^{(q)}$ is the residual matrix for block $q$ at transformed face $k$. Every least-squares deflation loading is evaluated on the selected residual face and divided by the squared norm of its regressor score. For a subject-level response, we define $\mathbf{Z}_{h,k}^{(2)} = \mathbf{R}_h$ at every candidate face. This shared response residual is updated once per component rather than once per face.

**Algorithm 2** Detailed two-block tPLS component extraction with canonical or regression deflation

**Input:** centred first-block tensor $\boldsymbol{X}^{(1)} \in \mathbb{R}^{n \times p_1 \times t}$; centred second longitudinal block $\boldsymbol{X}^{(2)} \in \mathbb{R}^{n \times p_2 \times t}$ or subject-level response $\mathbf{R}_1 \in \mathbb{R}^{n \times p_2}$; temporal-basis matrix $\mathbf{M}$; number of components $H$; deflation scheme $d \in \{\text{canonical}, \text{regression}\}$

transform each longitudinal input by $\times_3 \mathbf{M}$ and initialise its residual tensor
**for** $h = 1, \ldots, H$ **do**
  **for** $k = 1, \ldots, t$ **do**
    $\mathbf{G}_{h,k} \leftarrow \mathbf{Z}_{h,k}^{(1)\top} \mathbf{Z}_{h,k}^{(2)}$
    $[\mathbf{U}_{h,k}, \mathbf{S}_{h,k}, \mathbf{V}_{h,k}] \leftarrow \operatorname{svd}(\mathbf{G}_{h,k})$
    compute $s_{h,k}$ by Eq. 7 and, when a response is designated, $s_{h,k}^F$ by Eq. 8
  **end for**
  $k_h \leftarrow \arg\max_k s_{h,k}$ (canonical) or $\arg\max_k s_{h,k}^F$ (regression)
  $\underline{a}_h^{(1)} \leftarrow \mathbf{U}_{h,k_h}[:,1]$; $\underline{a}_h^{(2)} \leftarrow \mathbf{V}_{h,k_h}[:,1]$
  $\underline{b}_h^{(q)} \leftarrow \mathbf{Z}_{h,k_h}^{(q)} \underline{a}_h^{(q)}$, for $q = 1, 2$
  $\underline{p}_h^{(1)} \leftarrow \dfrac{\mathbf{Z}_{h,k_h}^{(1)\top} \underline{b}_h^{(1)}}{\underline{b}_h^{(1)\top} \underline{b}_h^{(1)}}$
  $\mathbf{D}_h^{(1)} \leftarrow \underline{b}_h^{(1)} \underline{p}_h^{(1)\top}$
  **if** $d =$ canonical **then**
    $\underline{p}_h^{(2)} \leftarrow \dfrac{\mathbf{Z}_{h,k_h}^{(2)\top} \underline{b}_h^{(2)}}{\underline{b}_h^{(2)\top} \underline{b}_h^{(2)}}$
    $\mathbf{D}_h^{(2)} \leftarrow \underline{b}_h^{(2)} \underline{p}_h^{(2)\top}$
  **else**
    $\underline{c}_h \leftarrow \dfrac{\mathbf{Z}_{h,k_h}^{(2)\top} \underline{b}_h^{(1)}}{\underline{b}_h^{(1)\top} \underline{b}_h^{(1)}}$
    $\mathbf{D}_h^{(2)} \leftarrow \underline{b}_h^{(1)} \underline{c}_h^{\top}$
  **end if**
  $\mathbf{Z}_{h+1,k_h}^{(1)} \leftarrow \mathbf{Z}_{h,k_h}^{(1)} - \mathbf{D}_h^{(1)}$
  **if** block 2 is longitudinal **then**
    $\mathbf{Z}_{h+1,k_h}^{(2)} \leftarrow \mathbf{Z}_{h,k_h}^{(2)} - \mathbf{D}_h^{(2)}$
    carry all unselected faces forward unchanged for both longitudinal blocks
  **else**
    $\mathbf{R}_{h+1} \leftarrow \mathbf{R}_h - \mathbf{D}_h^{(2)}$ and set $\mathbf{Z}_{h+1,k}^{(2)} = \mathbf{R}_{h+1}$ for every $k$
  **end if**
  for each longitudinal block, place $\mathbf{D}_h^{(q)}$ in face $k_h$ of an otherwise zero tensor and apply $\times_3 \mathbf{M}^{-1}$ to obtain its visit-domain component contribution
**end for**

**Output:** selected faces, loadings, scores, deflation loadings, response coefficients and visit-domain component contributions

### Appendix B Simulation models and evaluation protocol

#### B.1 Simulation A: single-omic data

Each simulated dataset comprised one omics assay, several visits per subject and two outcome groups. For subject $i$, feature $j$, visit $t$ and outcome group $g(i)$, the data were generated as

$$x_{ijt} = \mu_j + a_j z_{g(i)}(t) + b_{ij} + e_{ijt}. \tag{B.1}$$

where

- $\mu_j$ is the average level of feature $j$, shared by all subjects and visits.
- $z_g(t)$ is the true time course that distinguishes the two groups. The groups have the same average over time and differ only in trajectory shape (Section B.3), ensuring that the group signal is temporal rather than a difference in overall level.
- $a_j$ determines how strongly feature $j$ carries the group difference. For an informative feature, $a_j$ equals $+\delta$ or $-\delta$, with its sign chosen at random; for every other feature, $a_j = 0$. All informative features share $z_g(t)$, giving a rank-one group signal with a single underlying temporal pattern.
- $b_{ij}$ is a feature-specific, time-constant offset for subject $i$, drawn independently for each subject–feature pair. It contributes feature-specific rather than shared low-rank variation, although its magnitude can dominate the variance of an individual feature. Its constancy across visits places it entirely in the constant term of any temporal basis (Section B.3), where an estimator may mistake it for group signal.
- $e_{ijt}$ is residual noise, generated either independently or with an AR(1) correlation across neighbouring visits.

Consequently, group discrimination depends entirely on the shape of each feature's trajectory because the group-defining term $z_g(t)$ averages to zero over time.

#### B.2 Simulations B and C: multi-omics

Simulations B and C included multiple omics assays (blocks), indexed by $b$. For assay $b$, subject $i$, feature $j$ and visit $t$, the data were generated as

$$x_{ijt}^{(b)} = \mu_j^{(b)} + a_j^{(b)} z_{b,g(i)}(t) + s_{ij}^{(b)} + e_{ijt}^{(b)}. \tag{B.2}$$

The first two terms have the same roles as in Equation B.1: a feature mean and a group-discriminating temporal loading. The subject effect $s_{ij}^{(b)}$ is a time-constant offset for subject $i$ on feature $j$ of assay $b$, and determines whether stable subject-level variation is shared with other features or remains feature-specific. Each feature of assay $b$ is either *correlated* ($j \in \mathcal{C}_b$) or *uncorrelated* ($j \notin \mathcal{C}_b$). Correlated features load on a common subject factor, so their offsets covary across subjects:

$$s_{ij}^{(b)} = \begin{cases} \rho_b h_i + \sqrt{\sigma_b^2 - \rho_b^2}\, u_{ij}^{(b)}, & j \in \mathcal{C}_b \quad \text{(correlated)}, \\ \sigma_b u_{ij}^{(b)}, & j \notin \mathcal{C}_b \quad \text{(uncorrelated)}. \end{cases} \tag{B.3}$$

- $h_i \sim \mathcal{N}(0,1)$ is a subject-specific value shared by the correlated features of every assay.
- $u_{ij}^{(b)} \sim \mathcal{N}(0,1)$ is generated independently for every subject–feature pair.

**Table B.1. Feature classes defined by two independent properties.**

| Class | Discriminates groups? | Shares subject factor $h_i$? |
|---|---|---|
| CD (correlated, discriminative) | yes | yes |
| UD (uncorrelated, discriminative) | yes | no |
| CN (correlated, non-discriminative) | no | yes |
| UN (uncorrelated, non-discriminative) | no | no |

- $\sigma_b$ is the standard deviation of the offset. It is the same for correlated and uncorrelated features, so the two cases differ in correlation, not in scale.
- $0 \le \rho_b \le \sigma_b$ controls how strongly a correlated feature of assay $b$ follows $h_i$. Two correlated features, in assays $b$ and $b'$, have offset correlation $\rho_b\rho_{b'}/(\sigma_b\sigma_{b'})$, which is $(\rho_b/\sigma_b)^2$ within one assay and $\rho_b\rho_{b'}$ when $\sigma_b = \sigma_{b'} = 1$. Uncorrelated features are independent of all others.

Here, "correlated" refers specifically to the subject offset, and hence to correlation across subjects. A subject with a high $h_i$ tends to have high values for correlated features in every assay and at every visit because these features share the same underlying subject value. As this association is static, changes in cross-assay relationships over time arise from the temporal term $z_{b,g(i)}(t)$. Simulation B used this distinction to assess whether a fitted cross-assay correlation captured temporal coordination or static sharing through $s_{ij}^{(b)}$.

Each assay was also divided into four feature classes that crossed two independent properties: discrimination between groups and sharing of the subject factor $h_i$ (Table B.1 and Fig. B.1). The classes were correlated and discriminative (CD), uncorrelated and discriminative (UD), correlated and non-discriminative (CN), and uncorrelated and non-discriminative (UN).

The two properties were independent by construction. A UD feature discriminated between groups without sharing variation across assays, whereas a CN feature shared a subject-level tendency across assays without discriminating between groups. This separation allowed true-marker recovery and the selection of shared nuisance variation to be evaluated independently.

#### B.3 Temporal contrasts

The feature classes in Table B.1 determine which features carry a group difference, whereas the temporal contrast determines how that difference unfolds over the visits. The contrast corresponds to $z_g(t)$ in Eq. (B.1) and $z_{b,g}(t)$ for assay $b$ in Eq. (B.2). Every contrast gives the two groups the same average over six visits, making trajectory shape the only source of group discrimination. Linear and quadratic contrasts were used in Simulations A and B and as an anchor in Simulation C. Simulation B also included two contrasts whose energy was distributed across several temporal patterns, while Simulation C used rotated contrasts to induce disagreement among assays.

**Linear and quadratic contrasts** (Simulations A and B, and as an anchor in Simulation C). The groups had the same average over time and differed in whether their trajectories were linear or curved (Fig. B.2). The two shapes were orthogonal and scaled to the same length, so a given $\delta$ represented the same total signal strength for both.

**Off-basis-smooth and transient contrasts** (Simulation B, evaluating the temporal basis). For visit $u = 1, \ldots, 6$,

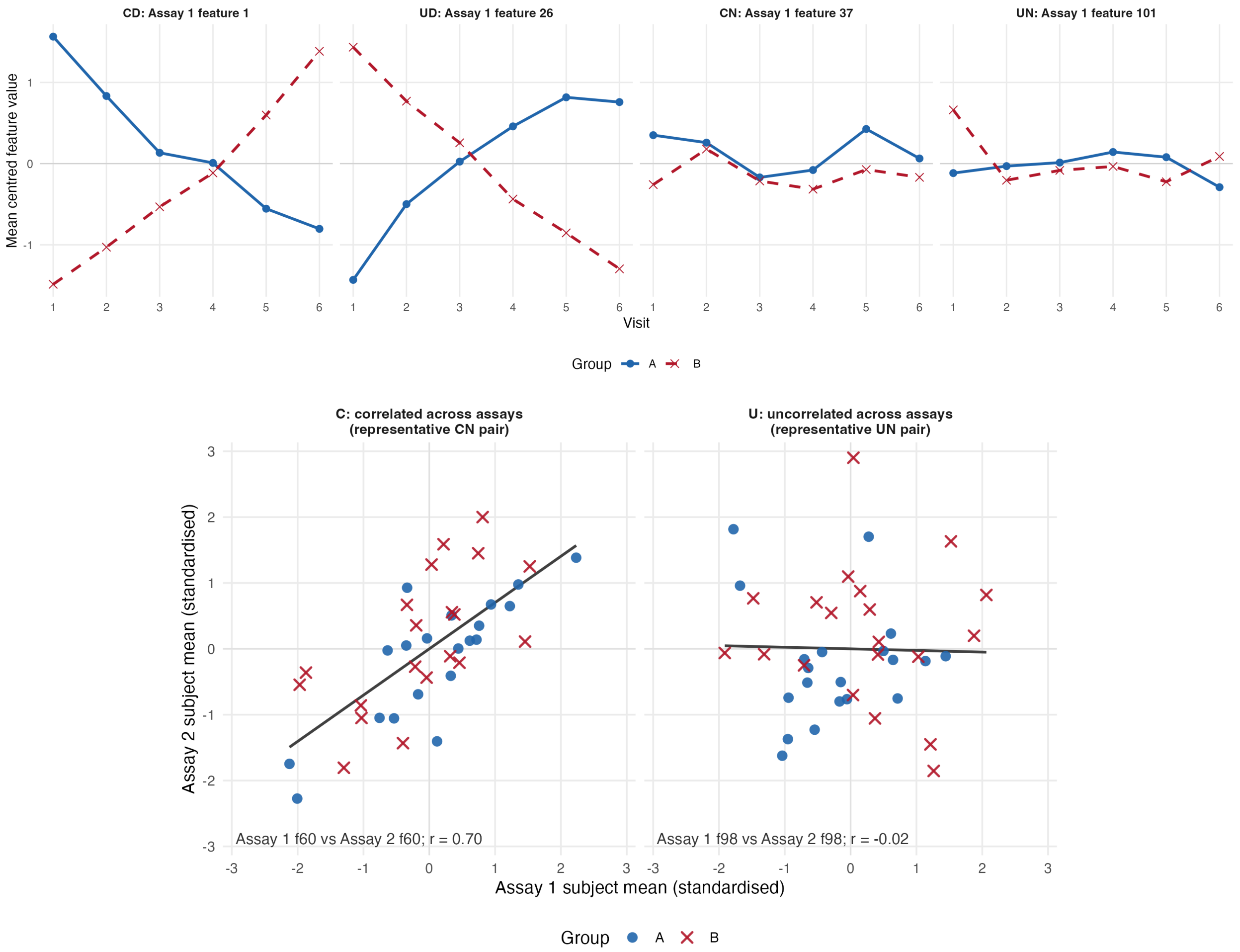


**Figure B.1. Feature classes in one simulated dataset.** The dataset contains 20 subjects per group, six visits and three assays, with a linear trajectory and $\delta = 0.5$. Top: group means across visits for one Assay 1 feature from each class. Correlated discriminative (CD) and uncorrelated discriminative (UD) features separate the groups; correlated non-discriminative (CN) and uncorrelated non-discriminative (UN) features show sampling variation only. Bottom: subject means, averaged across visits and standardised within each assay, for a matched feature pair in Assays 1 and 2. Correlated features share the subject factor $h_i$ ($r = 0.70$ in this example), whereas the displayed uncorrelated pair has $r = -0.02$.

$$g_{\text{smooth}}(u) = 0.60 \cos\left(\frac{\pi(u-0.5)}{6}\right) + 0.55 \cos\left(\frac{2\pi(u-0.5)}{6}\right) + 0.45 \cos\left(\frac{3\pi(u-0.5)}{6}\right), \tag{B.4}$$

$$g_{\text{transient}}(u) = \exp\left[-\frac{(u-4)^2}{2(0.6)^2}\right]. \tag{B.5}$$

The first contrast is a smooth mixture of three cosine patterns, with no single DCT basis direction containing more than 41.6% of its energy. The second is a short-lived Gaussian-shaped peak centred on visit 4 with a spread of approximately 0.6 visits; no single basis direction contains more than 52.0% of its energy. Both shapes were centred to zero mean and rescaled to the length of the linear contrast. The groups received mirror-image versions of each shape, preserving a common average and the interpretation of $\delta$ across all four contrasts. These patterns tested the temporal-basis representation when the true signal was distributed across several basis directions.

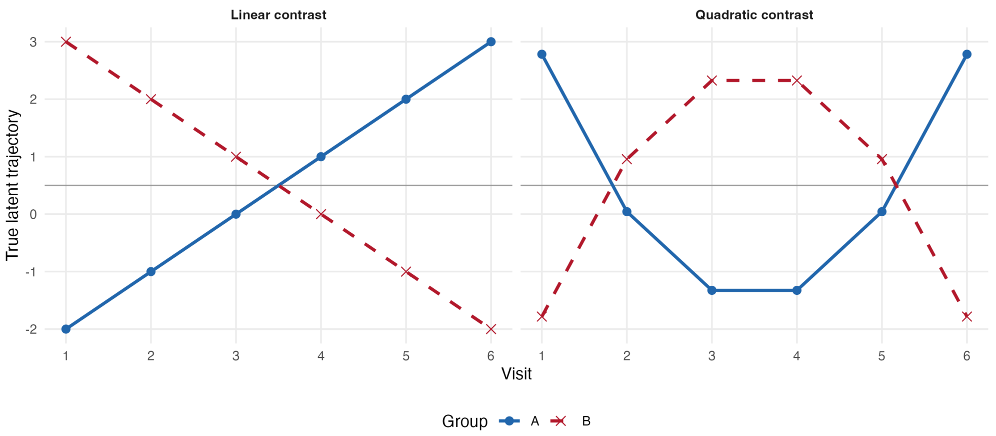


**Figure B.2. Linear and quadratic contrasts used in Simulation A.** Both groups have the same average across the six visits (grey line, 0.5) and differ only in trajectory shape.

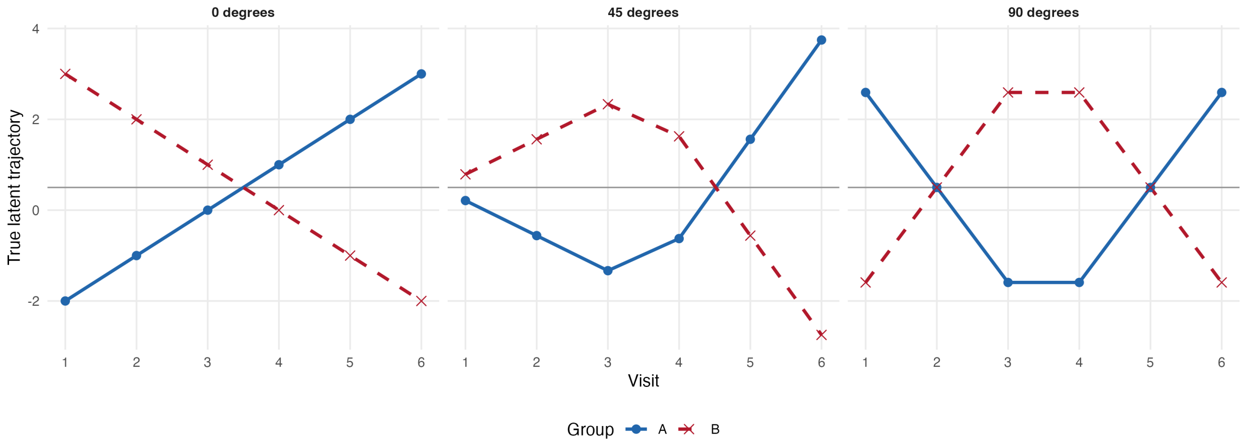


**Figure B.3. Rotated temporal contrasts used in Simulation C.** Omics 1 retains the linear contrast at every angle; the contrasts for omics 2 and 3 are shown at rotations of 0°, 45° and 90°. All panels have the same group time-average and contrast length.

**Rotated contrasts** (Simulation C, evaluating temporal disagreement among assays). Assay 1 retained the linear contrast, while Assays 2 and 3 used

$$z_b = \cos(\theta_b)\, z_{\text{linear}} + \sin(\theta_b)\, q, \tag{B.6}$$

where $q$ is a fixed, zero-mean shape with the same length as, and orthogonal to, the linear contrast. As $\theta_b$ increased from 0° to 90°, the patterns in assays 2 and 3 rotated away from the linear pattern in assay 1 without changing their total signal. All assays shared the same pattern at 0°, whereas the patterns in assays 2 and 3 were orthogonal to that in assay 1 at 90°. The rotation preserved the effect size, and the data generated for assay 1 were identical at every angle.

#### B.4 Simulation grids (Simulations A–C)

Table B.2 lists the settings of the three simulation designs. Error rates use 5-fold whole-subject cross-validation repeated 10 times.

**Table B.2. Simulation grids for Simulation A (single block), Simulation B (representation and basis scaling) and Simulation C (multiblock temporal disagreement).** "Cells" are the distinct parameter combinations evaluated.

| Parameter | Values tested |
|---|---|
| **A. Single-block temporal-shape recovery (Simulation A)** | |
| Assay | 1 assay, 200 features (15 informative), 6 visits |
| Subjects per group | 10, 20, 50 |
| $\delta$ | 0.05, 0.10, 0.15, 0.20, 0.30 |
| Subject-effect SD | 0, 0.5, 1, 2 |
| Residual AR(1) correlation | 0, 0.5 |
| Temporal contrast | linear, quadratic |
| Fixed | residual SD 1; shared subject factor off; 3 components |
| Methods | tPLS-DA; PLS-DA on flattened visits; tPCA; PCA on flattened visits; N-PLS; TEMPTED (centred, uncentred); PARAFAC; RM-ASCA+ |
| Cells × datasets | 22 × 50 |
| **B. Representation, compression and basis scaling (Simulation B)** | |
| Assays | 3 assays, 200, 150, 100 features (all with signal), 6 visits |
| $\delta$ | 0.1, 0.3 |
| Residual AR(1) correlation | 0, 0.6 |
| Cross-assay correlation scale | 0, 0.5 |
| Temporal contrast | linear, quadratic, off-basis-smooth, transient |
| DCT-compressed matrix | $k = 2, 3, 6$ coefficients retained |
| Design weight | 0 |
| Scaling diagnostic | $\delta = 0.1$, linear contrast, column scaling on vs. off |
| Fixed | 20 per group; subject-effect and residual SD 1; 3 components; top 30 features |
| Methods | block.tPLS-DA; block PLS-DA on flattened visits; DIABLO |
| Cells × datasets | 32 × 10 |
| **C. Multiblock temporal disagreement (Simulation C)** | |
| Assays | 3 assays, 200, 150, 100 features (all with signal), 6 visits |
| Rotation angle (assays 2, 3 vs. assay 1) | $0°, 30°, 45°, 60°, 75°, 90°$ |
| $\delta$ | 0.1, 0.2, 0.3 |
| Design weight | 0, 0.2 |
| Fixed | linear anchor contrast; 20 per group; full static cross-assay correlation; subject-effect and residual SD 1; independent residuals; 3 components; top 30 features |
| Methods | block.tPLS-DA; block PLS-DA on flattened visits; DIABLO |
| Cells × datasets | 36 × 50 |

Simulation A varied one factor at a time around a reference setting of 20 subjects per group, $\delta = 0.2$, subject-effect SD 1, residual SD 1 and independent residuals for each contrast, rather than crossing all values. Simulation B used 10 datasets per cell, and scaling sensitivity was evaluated in a focused diagnostic rather than for every matrix comparator. The main text therefore reports the mechanism underlying the representation limitation and the equality observed under matched scaling. In Simulation C, Assays 2 and 3 rotated together, creating a two-against-one temporal pattern. The results describe this rotation scheme, while selection stability and a general threshold were outside the scope of the simulation.

### Appendix C Real-data cohorts

We analysed three longitudinal multi-omics cohorts, each with two omic types (Table 2). The cohort sizes in Table 2were 90 of the 171 CMI-PB subjects (aP 44, wP 46), 52 of 80 HE INFANTS infants (breast-fed 9, standard formula 23, MFGM formula 20) and 46 of 75 MARS subjects (healthy 14, IBS-C 11, IBS-D 21).

Each omic block was pre-processed once, for every method, so that all methods were fitted on identical inputs. In CMI-PB, the 31 antibody titres are already expressed as ratios to the baseline median and were $\log_2$ transformed; the gene-expression counts had been batch-corrected using ComBat-seq by the original authors (Zhang et al., 2020), , and we retained the 1,000 most variable of 6,650 genes before applying TMM normalisation and a $\log_2$-CPM transformation. In HE INFANTS, 124 of 295 genera passed a low-abundance filter and were centred log-ratio transformed, and 118 of 120 NMR metabolites were $\log_2$ transformed. In MARS, event-triggered flare samples were excluded, 301 of 2,895 genera passed a pre-filter at 0.01% of the total counts and were total-sum scaled then centred log-ratio transformed, and 26 of 43 faecal metabolites were retained, with values below the limit of detection replaced by half the smallest positive value of that metabolite before the $\log_2$ transformation.

Two further steps were needed for the methods to be comparable. First, the HE INFANTS metabolome included 34.4% zeros, and 17 of its 118 metabolites were below detection in every subject at at least one visit. Thus, some features had zero variance within a fold. We therefore only considered 101 metabolites. Second, TEMPTED and PARAFAC apply a centred log-ratio transformation internally by default; since every block here is already transformed, that option was disabled.

### Appendix D Additional benchmark results

We report the numerical results underlying the simulation and real-data benchmarks. Unless stated otherwise, balanced error rates (BERs) are the mean (SD) across the specified cross-validation repeats; lower values indicate better classification, and chance is approximately 0.50 for two groups. Bold identifies the best value within each method family, with supervised and unsupervised methods treated as separate families. PLS-DA on flattened visits represents each feature–visit combination as a separate variable and does not encode visit order. Rows labelled "without face scaling" use the raw covariance score for face selection instead of the scaled score in Section 2.3.3; all other settings match those of tPLS-DA. Refer to Section 2.6.4 regarding how the unsupervised and supervised methods were evaluated.

**Table D.1. Simulation A (Single-omic simulation): BER by effect size for linear and quadratic temporal contrasts.** Top: linear contrast. Bottom: quadratic contrast. Each dataset contains 200 features (15 informative), six visits and 20 subjects per group, with between-subject SD 1 and residual SD 1; each column summarises 50 datasets. RM-ASCA+ evaluates a population-level interaction and is therefore omitted from the individual-level BER table.

| Linear temporal contrast | | | | | |
|---|---|---|---|---|---|
| Method | $\delta = 0.05$ | $\delta = 0.1$ | $\delta = 0.15$ | $\delta = 0.2$ | $\delta = 0.3$ |
| tPLS-DA | **0.450 (0.085)** | **0.229 (0.074)** | **0.068 (0.039)** | **0.010 (0.012)** | **0.000 (0.000)** |
| tPLS-DA without face scaling | 0.502 (0.068) | 0.502 (0.068) | 0.502 (0.068) | 0.502 (0.068) | 0.502 (0.068) |
| PLS-DA on flattened visits | 0.484 (0.068) | 0.441 (0.068) | 0.370 (0.068) | 0.278 (0.064) | 0.094 (0.042) |
| N-PLS | 0.501 (0.068) | 0.501 (0.068) | 0.501 (0.069) | 0.501 (0.069) | 0.491 (0.066) |
| ***Unsupervised methods*** *(separate comparison family)* | | | | | |
| tPCA | 0.518 (0.068) | 0.518 (0.068) | 0.518 (0.068) | 0.518 (0.068) | 0.518 (0.068) |
| PCA on flattened visits | 0.515 (0.068) | **0.504 (0.069)** | **0.480 (0.071)** | **0.433 (0.074)** | **0.223 (0.071)** |
| TEMPTED | 0.515 (0.066) | 0.516 (0.066) | 0.515 (0.066) | 0.515 (0.066) | 0.513 (0.064) |
| TEMPTED, uncentred | **0.509 (0.061)** | 0.509 (0.061) | 0.508 (0.061) | 0.508 (0.060) | 0.508 (0.059) |
| PARAFAC (transductive) | 0.511 (0.081) | 0.507 (0.081) | 0.506 (0.080) | 0.506 (0.082) | 0.504 (0.089) |
| Quadratic temporal contrast | | | | | |
| Method | $\delta = 0.05$ | $\delta = 0.1$ | $\delta = 0.15$ | $\delta = 0.2$ | $\delta = 0.3$ |
| tPLS-DA | **0.461 (0.077)** | **0.215 (0.064)** | **0.061 (0.037)** | **0.010 (0.013)** | **0.000 (0.000)** |
| tPLS-DA without face scaling | 0.502 (0.068) | 0.502 (0.068) | 0.502 (0.068) | 0.502 (0.068) | 0.502 (0.068) |
| PLS-DA on flattened visits | 0.482 (0.069) | 0.438 (0.069) | 0.363 (0.067) | 0.271 (0.062) | 0.089 (0.037) |
| N-PLS | 0.501 (0.069) | 0.501 (0.069) | 0.501 (0.069) | 0.501 (0.069) | 0.489 (0.065) |
| ***Unsupervised methods*** *(separate comparison family)* | | | | | |
| tPCA | 0.518 (0.068) | 0.518 (0.068) | 0.518 (0.068) | 0.518 (0.068) | 0.518 (0.068) |
| PCA on flattened visits | 0.516 (0.070) | **0.506 (0.073)** | **0.482 (0.075)** | **0.436 (0.078)** | **0.223 (0.074)** |
| TEMPTED | 0.516 (0.066) | 0.515 (0.066) | 0.515 (0.066) | 0.514 (0.066) | 0.514 (0.066) |
| TEMPTED, uncentred | **0.509 (0.062)** | 0.509 (0.062) | 0.509 (0.062) | 0.509 (0.062) | 0.509 (0.061) |
| PARAFAC (transductive) | 0.510 (0.080) | **0.506 (0.083)** | 0.505 (0.083) | 0.503 (0.086) | 0.499 (0.085) |

**Table D.2. Single-omic simulation (Simulation A): sensitivity to cohort size, between-subject variation and residual correlation.** The linear contrast uses $\delta = 0.2$, with one factor varied at a time from the reference setting of 20 subjects per group, between-subject SD 1 and independent residuals. Each column summarises 50 datasets; the reference column corresponds to the linear contrast at $\delta = 0.2$ in Table D.1.

| Method | 10 per group | reference | 50 per group | subject SD 0 | subject SD 0.5 | subject SD 2 | residual AR(1) 0.5 |
|---|---|---|---|---|---|---|---|
| tPLS-DA | **0.019 (0.023)** | **0.010 (0.012)** | **0.006 (0.006)** | **0.010 (0.011)** | **0.010 (0.012)** | **0.010 (0.013)** | **0.042 (0.026)** |
| tPLS-DA without face scaling | 0.496 (0.120) | 0.502 (0.068) | 0.496 (0.054) | **0.010 (0.010)** | 0.165 (0.145) | 0.498 (0.066) | 0.500 (0.081) |
| PLS-DA on flattened visits | 0.333 (0.116) | 0.278 (0.064) | 0.177 (0.042) | 0.072 (0.043) | 0.120 (0.048) | 0.426 (0.065) | 0.312 (0.070) |
| N-PLS | 0.494 (0.119) | 0.501 (0.069) | 0.479 (0.058) | 0.019 (0.015) | 0.259 (0.136) | 0.494 (0.073) | 0.498 (0.078) |
| ***Unsupervised methods*** *(separate comparison family)* | | | | | | | |
| tPCA | 0.479 (0.112) | 0.518 (0.068) | 0.491 (0.043) | **0.010 (0.011)** | 0.515 (0.062) | 0.512 (0.071) | 0.513 (0.064) |
| PCA on flattened visits | **0.403 (0.130)** | **0.433 (0.074)** | **0.394 (0.060)** | 0.100 (0.047) | **0.206 (0.071)** | **0.499 (0.076)** | **0.452 (0.074)** |
| TEMPTED | 0.480 (0.114) | 0.515 (0.066) | 0.495 (0.043) | 0.023 (0.023) | 0.386 (0.094) | 0.514 (0.070) | 0.512 (0.063) |
| TEMPTED, uncentred | 0.474 (0.110) | 0.508 (0.060) | 0.496 (0.042) | 0.033 (0.039) | 0.434 (0.089) | 0.507 (0.068) | 0.508 (0.061) |
| PARAFAC (transductive) | 0.487 (0.125) | 0.506 (0.082) | 0.491 (0.057) | 0.031 (0.027) | 0.494 (0.090) | 0.512 (0.094) | 0.508 (0.078) |

**Table D.3. Single-omic simulation (Simulation A): recovery of informative features and temporal contrasts.** Top: recall among the 15 informative features, false-positive rate among null features, and Jaccard overlap of selections across folds. Random-selection references are 0.075 for recall and false-positive rate and 0.039 for Jaccard overlap. Bottom: non-constancy of the fitted temporal loading, which is 0 for a loading that does not change over time and about 0.833 for a random loading at six visits, and its absolute correlation with the true group contrast. The empirical null values for agreement are 0.833 for the linear contrast and 0.854 for the quadratic contrast. Results summarise 50 datasets from the reference setting ($\delta = 0.2$); higher values are better except for the false-positive rate. Bold identifies the best value in each column within a method family. PARAFAC uses a joint, transductive fit to all subjects.

Feature selection.

| Method | recall, linear | recall, quadratic | false positives, linear | false positives, quadratic | Jaccard, linear | Jaccard, quadratic |
|---|---|---|---|---|---|---|
| tPLS-DA | **0.763 (0.071)** | **0.771 (0.068)** | **0.019 (0.006)** | **0.019 (0.005)** | **0.480 (0.061)** | **0.480 (0.056)** |
| tPLS-DA without face scaling | 0.069 (0.077) | 0.069 (0.077) | 0.075 (0.006) | 0.075 (0.006) | 0.143 (0.034) | 0.143 (0.034) |
| PLS-DA on flattened visits | 0.423 (0.100) | 0.383 (0.120) | 0.047 (0.008) | 0.050 (0.010) | 0.198 (0.047) | 0.179 (0.047) |
| N-PLS | 0.068 (0.079) | 0.065 (0.068) | 0.076 (0.006) | 0.076 (0.006) | 0.130 (0.035) | 0.131 (0.034) |
| ***Unsupervised methods*** *(separate comparison family)* | | | | | | |
| tPCA | 0.071 (0.062) | 0.071 (0.062) | 0.075 (0.005) | 0.075 (0.005) | 0.180 (0.039) | 0.180 (0.039) |
| PCA on flattened visits | **0.119 (0.081)** | **0.100 (0.081)** | **0.071 (0.007)** | **0.073 (0.007)** | 0.149 (0.039) | 0.149 (0.033) |
| TEMPTED | 0.069 (0.059) | 0.075 (0.060) | 0.075 (0.005) | 0.075 (0.005) | 0.182 (0.035) | 0.178 (0.035) |
| TEMPTED, uncentred | 0.063 (0.069) | 0.069 (0.071) | 0.076 (0.006) | 0.075 (0.006) | **0.315 (0.070)** | **0.312 (0.070)** |
| PARAFAC (transductive) | 0.052 (0.054) | 0.044 (0.051) | 0.077 (0.004) | 0.078 (0.004) | 0.117 (0.033) | 0.123 (0.029) |

Recovery of the temporal contrast.

| Method | non-constancy, linear | non-constancy, quadratic | agreement, linear | agreement, quadratic |
|---|---|---|---|---|
| tPLS-DA | **1.000 (0.000)** | **1.000 (0.000)** | **0.995 (0.000)** | **0.982 (0.000)** |
| N-PLS | 0.011 (0.012) | 0.010 (0.010) | 0.597 (0.314) | 0.636 (0.308) |
| TEMPTED | 0.002 (0.002) | 0.002 (0.002) | 0.382 (0.266) | 0.352 (0.223) |
| TEMPTED, uncentred | 0.000 (0.000) | 0.000 (0.000) | 0.381 (0.250) | 0.362 (0.214) |
| PARAFAC (transductive) | 0.003 (0.002) | 0.003 (0.002) | 0.374 (0.242) | 0.427 (0.253) |

*Recovery of the temporal contrast, Table D.3.* Methods that retain an explicit temporal mode return a sixelement temporal loading that characterises the pattern across visits contributing to group separation. For tPLS-DA, this loading is the selected discrete cosine transform (DCT) face; for N-PLS, TEMPTED and PARAFAC, it is estimated from the data. Flattened-matrix methods do not retain a separate temporal mode, so this measure is not applicable to them.

Temporal agreement was quantified as the absolute correlation between the estimated temporal loading and the true group contrast. With only six visits, an unrelated loading can correlate strongly with a smooth contrast by chance. Agreement was therefore evaluated against an empirical null threshold, defined as the 95th percentile of the correlations obtained from random loadings: 0.833 for the linear contrast and 0.854 for the quadratic contrast. Only tPLS-DA exceeded these thresholds, with agreement values of 0.995 and 0.982, respectively. N-PLS, TEMPTED and PARAFAC remained below the corresponding thresholds for both contrasts, with a maximum agreement of 0.636, and therefore did not meet the null-calibrated criterion for recovery of the temporal contrast.

Non-constancy provides a complementary measure of whether a temporal loading varies across visits, irrespective of whether that variation agrees with the true contrast. It should therefore be interpreted alongside temporal agreement. Non-constancy is 0 for a loading that is constant over time and approximately 0.833 for a random loading across six visits. Values of 0.011 or lower for N-PLS, TEMPTED and PARAFAC indicate that their estimated temporal loadings were essentially constant, consistent with separation driven primarily by differences in overall level rather than by changes over time.

**Table D.4. Simulation B (classification and temporal-contrast recovery for four trajectory shapes).** Top: balanced error rate. Bottom: absolute correlation between the fitted temporal loading and the true contrast. For a method restricted to one temporal direction per component, the maximum possible agreement is 0.645 for the off-basis-smooth contrast and 0.721 for the transient contrast. The setting contains three omics, $\delta = 0.1$, design weight 0, independent residuals and cross-omics correlation set to 0; each column summarises 10 datasets. The matrix-based methods use column scaling whereas block.tPLS-DA uses centred, unscaled data; Table D.5 isolates this scaling effect.

| Balanced error rate | | | | |
|---|---|---|---|---|
| Method | linear | quadratic | off-basis smooth | transient |
| block.tPLS-DA | **0.041 (0.027)** | **0.052 (0.042)** | **0.084 (0.034)** | **0.076 (0.054)** |
| block PLS-DA, flattened visits | 0.270 (0.075) | 0.279 (0.079) | 0.285 (0.074) | 0.278 (0.079) |
| DIABLO | 0.310 (0.079) | 0.350 (0.063) | 0.306 (0.076) | 0.305 (0.081) |
| **Agreement of the fitted temporal loading with the true contrast** | | | | |
| Method | linear | quadratic | off-basis smooth | transient |
| block.tPLS-DA | **0.995 (0.000)** | **0.982 (0.000)** | 0.634 (0.023) | 0.721 (0.000) |
| block PLS-DA, flattened visits | 0.444 (0.294) | 0.529 (0.282) | 0.385 (0.244) | 0.448 (0.309) |
| DIABLO | 0.709 (0.144) | 0.675 (0.259) | **0.944 (0.014)** | **0.842 (0.242)** |

**Table D.5. Simulation B (effect of column scaling on flattened and DCT-transformed data).** The same data were supplied to block PLS-DA as flattened visits or as all six DCT terms, which form an exact rotation of the flattened data ($\delta = 0.1$, linear contrast, five datasets, five-fold cross-validation repeated five times). The rows differ only in matrix-based method column scaling. With scaling switched off, the two representations gave the same balanced error rate in every dataset (mean difference 0.000, SD 0.000); with scaling on they differed by 0.195 in favour of the DCT. The gap is therefore produced by the scaling and not by the representation.

| matrix-based column scaling | flattened visits | all 6 DCT terms |
|---|---|---|
| on (default) | 0.297 (0.050) | 0.102 (0.031) |
| off | 0.310 (0.042) | 0.310 (0.042) |

**Table D.6. Simulation C: BER across temporal-disagreement angles at design weights 0 and 0.2.** Top: design weight 0. Bottom: design weight 0.2. Each cell summarises 50 datasets; bold identifies the best value in each column within each $\delta$. Block PLS-DA and DIABLO use the default column scaling (Table D.5).

| $\delta$ | Method | 0° | 30° | 45° | 60° | 75° | 90° |
|---|---|---|---|---|---|---|---|
| Design weight 0 | | | | | | | |
| 0.1 | block.tPLS-DA | **0.138 (0.088)** | **0.142 (0.087)** | **0.146 (0.089)** | **0.154 (0.079)** | **0.138 (0.073)** | **0.135 (0.074)** |
| 0.1 | block PLS-DA, flattened visits | 0.309 (0.062) | 0.314 (0.061) | 0.315 (0.062) | 0.314 (0.062) | 0.315 (0.063) | 0.314 (0.063) |
| 0.1 | DIABLO | 0.348 (0.083) | 0.344 (0.080) | 0.339 (0.080) | 0.347 (0.080) | 0.359 (0.083) | 0.367 (0.085) |
| 0.2 | block.tPLS-DA | **0.031 (0.039)** | **0.042 (0.047)** | 0.050 (0.060) | **0.043 (0.055)** | **0.051 (0.048)** | **0.042 (0.039)** |
| 0.2 | block PLS-DA, flattened visits | 0.058 (0.031) | 0.061 (0.032) | 0.062 (0.033) | 0.060 (0.032) | 0.057 (0.031) | 0.056 (0.031) |
| 0.2 | DIABLO | 0.057 (0.030) | **0.042 (0.022)** | **0.038 (0.020)** | 0.044 (0.024) | 0.062 (0.029) | 0.071 (0.030) |
| 0.3 | block.tPLS-DA | 0.004 (0.009) | 0.006 (0.013) | 0.011 (0.020) | 0.012 (0.025) | 0.012 (0.017) | 0.007 (0.012) |
| 0.3 | block PLS-DA, flattened visits | 0.004 (0.007) | 0.005 (0.008) | 0.005 (0.008) | 0.004 (0.008) | 0.004 (0.007) | 0.004 (0.007) |
| 0.3 | DIABLO | **0.001 (0.002)** | **0.000 (0.001)** | **0.000 (0.000)** | **0.000 (0.001)** | **0.001 (0.001)** | **0.002 (0.002)** |
| Design weight 0.2 | | | | | | | |
| $\delta$ | Method | 0° | 30° | 45° | 60° | 75° | 90° |
| 0.1 | block.tPLS-DA | 0.471 (0.087) | 0.471 (0.087) | 0.471 (0.087) | 0.471 (0.087) | 0.471 (0.088) | 0.470 (0.088) |
| 0.1 | block PLS-DA, flattened visits | **0.316 (0.058)** | **0.320 (0.060)** | 0.321 (0.060) | 0.321 (0.061) | **0.320 (0.061)** | **0.319 (0.061)** |
| 0.1 | DIABLO | 0.321 (0.071) | 0.321 (0.066) | **0.315 (0.064)** | **0.316 (0.061)** | 0.322 (0.063) | 0.324 (0.067) |
| 0.2 | block.tPLS-DA | 0.166 (0.054) | 0.197 (0.049) | 0.237 (0.046) | 0.361 (0.086) | 0.268 (0.045) | 0.257 (0.046) |
| 0.2 | block PLS-DA, flattened visits | 0.061 (0.031) | 0.065 (0.031) | 0.065 (0.032) | 0.063 (0.031) | 0.062 (0.032) | 0.060 (0.031) |
| 0.2 | DIABLO | **0.037 (0.020)** | **0.027 (0.015)** | **0.025 (0.013)** | **0.028 (0.014)** | **0.035 (0.015)** | **0.040 (0.019)** |
| 0.3 | block.tPLS-DA | 0.042 (0.041) | 0.066 (0.049) | 0.094 (0.052) | 0.070 (0.046) | 0.150 (0.054) | 0.139 (0.054) |
| 0.3 | block PLS-DA, flattened visits | 0.004 (0.007) | 0.005 (0.008) | 0.005 (0.008) | 0.005 (0.008) | 0.004 (0.007) | 0.004 (0.007) |
| 0.3 | DIABLO | **0.000 (0.001)** | **0.000 (0.000)** | **0.000 (0.000)** | **0.000 (0.000)** | **0.000 (0.001)** | **0.001 (0.002)** |

**Table D.7. Simulation C: recovery of UD features and block-specific temporal patterns at design weight 0 and $\delta = 0.3$.** Top: uncorrelated discriminative (UD) feature recall averaged equally across the three omics. Bottom: block-specific UD recall, agreement between each fitted temporal loading and the true contrast for that omic, and the face selected for Component 1. Omics 1 retains the linear contrast at every angle, while omics 2 and 3 are rotated. Results summarise 50 datasets.

UD feature recall averaged equally across the three omics. (higher values are better; chance is 0.217)

| Method | 0° | 30° | 45° | 60° | 75° | 90° |
|---|---|---|---|---|---|---|
| block.tPLS-DA | **0.698 (0.050)** | **0.751 (0.078)** | **0.793 (0.081)** | **0.787 (0.072)** | **0.697 (0.077)** | **0.624 (0.063)** |
| block PLS-DA, flattened visits | 0.622 (0.093) | 0.663 (0.094) | 0.675 (0.099) | 0.660 (0.103) | 0.613 (0.106) | 0.593 (0.094) |
| DIABLO | 0.463 (0.070) | 0.478 (0.072) | 0.480 (0.071) | 0.467 (0.069) | 0.465 (0.078) | 0.446 (0.074) |

block.tPLS-DA by omics (Higher values indicate better recovery; the chance references for UD recall are 0.150 for omics 1 and 0.250 averaged across omics 2 and 3).

| angle | UD recall, omics 1 | UD recall, omics 2–3 | time-shape agreement, omics 1 | time-shape agreement, omics 2–3 | face chosen by component 1 |
|---|---|---|---|---|---|
| 0° | 0.767 (0.091) | 0.664 (0.077) | 0.995 (0.000) | 0.995 (0.000) | face 2 in 50 |
| 30° | 0.767 (0.093) | 0.743 (0.114) | 0.995 (0.000) | 0.862 (0.000) | face 2 in 50 |
| 45° | 0.775 (0.074) | 0.803 (0.125) | 0.995 (0.000) | 0.704 (0.000) | face 2 in 50 |
| 60° | 0.760 (0.102) | 0.801 (0.101) | 0.836 (0.368) | 0.556 (0.136) | face 2 in 42, face 3 in 8 |
| 75° | 0.628 (0.129) | 0.731 (0.079) | 0.000 (0.000) | 0.966 (0.000) | face 3 in 50 |
| 90° | 0.553 (0.101) | 0.659 (0.082) | 0.000 (0.000) | 1.000 (0.000) | face 3 in 50 |

**Table D.8. Longitudinal cohorts: sensitivity of supervised classification to feature scaling.** We report the balanced error rate (BER), mean (SD) across 10 repeats of five-fold subject-level cross-validation when features were scaled, and the difference from unscaled features that were already reported in Table 5 (scaled minus unscaled; negative values indicate lower error after scaling). All models used three components, and scaling parameters were estimated within each training fold. Chance BER was 0.500 for CMI-PB and 0.667 for HE INFANTS and MARS. Across the six omics individual blocks, scaling changed the method with the lowest BER in three, but no method was consistently best. Bold identifies the lowest scaled BER in each block.

| Method | CMI-PB ($n = 90$) | | HE INFANTS ($n = 52$) | | MARS ($n = 46$) | |
|---|---|---|---|---|---|---|
| | antibody titres | gene expression | genera | metabolome | genera | metabolome |
| tPLS-DA | 0.362 (0.035) (+0.053) | 0.404 (0.040) (+0.009) | 0.574 (0.052) (−0.026) | 0.483 (0.057) (+0.076) | 0.567 (0.047) (+0.008) | 0.510 (0.055) (+0.101) |
| N-PLS | **0.302 (0.027)** (−0.019) | 0.371 (0.018) (−0.001) | **0.426 (0.061)** (−0.063) | **0.362 (0.028)** (−0.025) | 0.654 (0.058) (+0.005) | 0.636 (0.036) (+0.090) |
| PLS-DA on flattened visits | 0.320 (0.033) (+0.002) | **0.311 (0.025)** (−0.037) | 0.514 (0.053) (−0.061) | 0.413 (0.028) (+0.047) | **0.542 (0.036)** (−0.016) | **0.383 (0.034)** (−0.123) |

**Table D.9. Longitudinal cohorts: stability of feature selection by the supervised methods.** Stability is the Jaccard similarity between the top 30 features selected in different folds; parentheses give the ratio to the expected overlap of two random sets of 30 features. CMI-PB antibody titres and the MARS metabolome are omitted because 30 of 31 and 26 of 26 features, respectively, would be selected. Across the four blocks assessed, overlap exceeded the random-set reference for every method and scaling condition, but no method was consistently the most stable. tPLS-DA had the lowest overlap in three of four unscaled blocks and all four scaled blocks. Higher values indicate greater reproducibility; bold identifies the largest similarity in each column within a block.

CMI-PB, gene expression: 1000 features, top 30 selected, random-set reference 0.0152

| Method | Jaccard, unscaled (ratio to reference) | Jaccard, scaled (ratio to reference) |
|---|---|---|
| tPLS-DA | 0.398 (26.2×) | 0.134 (8.8×) |
| PLS-DA on flattened visits | 0.365 (24.0×) | **0.203** (13.3×) |
| N-PLS | **0.430** (28.3×) | 0.154 (10.1×) |

HE INFANTS, genera: 124 features, top 30 selected, random-set reference 0.1376

| Method | Jaccard, unscaled (ratio to reference) | Jaccard, scaled (ratio to reference) |
|---|---|---|
| tPLS-DA | 0.480 (3.5×) | 0.324 (2.4×) |
| PLS-DA on flattened visits | **0.614** (4.5×) | 0.471 (3.4×) |
| N-PLS | 0.571 (4.1×) | **0.508** (3.7×) |

HE INFANTS, metabolome: 101 features, top 30 selected, random-set reference 0.1744

| Method | Jaccard, unscaled (ratio to reference) | Jaccard, scaled (ratio to reference) |
|---|---|---|
| tPLS-DA | 0.465 (2.7×) | 0.387 (2.2×) |
| PLS-DA on flattened visits | **0.553** (3.2×) | **0.432** (2.5×) |
| N-PLS | 0.504 (2.9×) | 0.426 (2.4×) |

MARS, genera: 301 features, top 30 selected, random-set reference 0.0524

| Method | Jaccard, unscaled (ratio to reference) | Jaccard, scaled (ratio to reference) |
|---|---|---|
| tPLS-DA | 0.274 (5.2×) | 0.123 (2.4×) |
| PLS-DA on flattened visits | 0.312 (5.9×) | 0.175 (3.3×) |
| N-PLS | **0.325** (6.2×) | **0.221** (4.2×) |

### Appendix E Batch correction and design-weight sensitivity for the FMT study

Exploratory analyses identified an unrecorded systematic source of variation in the mOTU and KO data blocks. The pattern was first detected by tPCA (Fig. E.1**A**) and confirmed by separate PCA analyses at each time point (Fig. E.1**B**). As laboratory batch records were unavailable, we defined two *in silico* batches by separating subjects on the tPCA sample plot. For all 21 subjects, this separation coincided with the sign of the first tPCA component in the uncorrected mOTU data. The systematic variation was consistent with a batch effect that could confound the biological signal and reduce power to detect treatment differences.

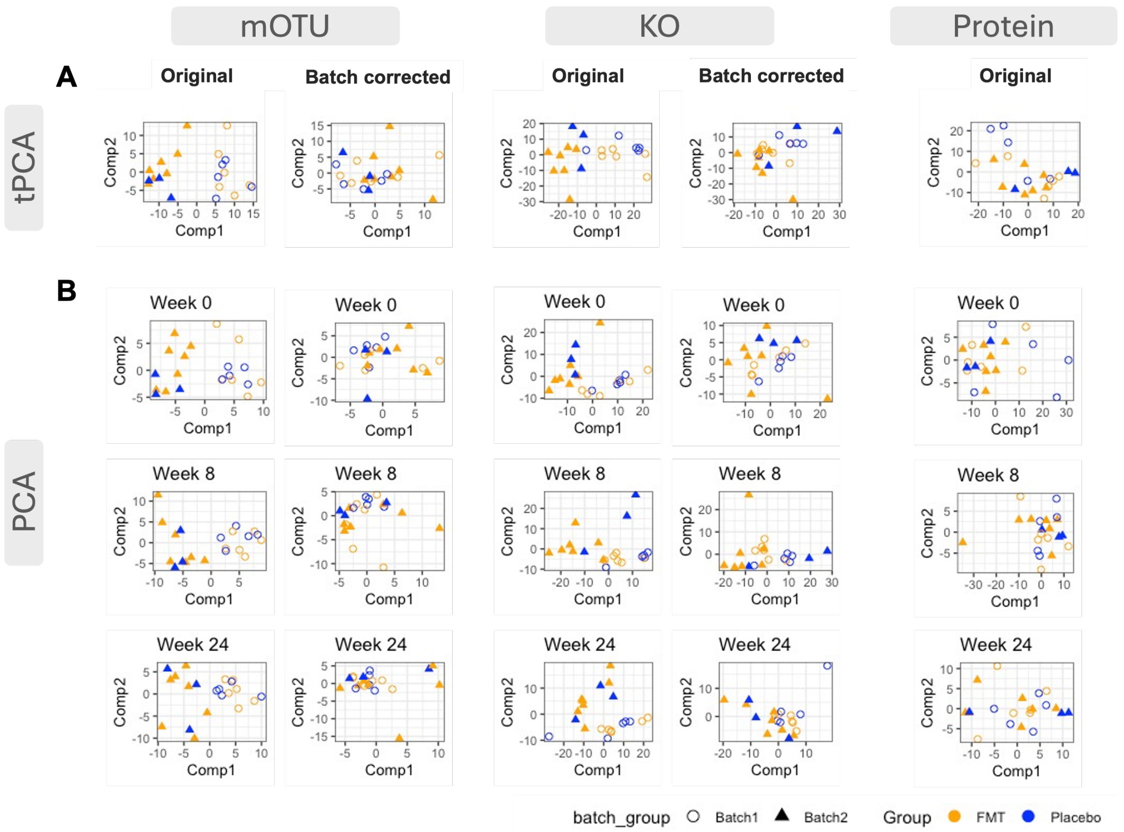


**Figure E.1. Detection and correction of systematic variation in the fecal microbiota transplantation study. A**: tensor principal component analysis (tPCA) sample plots before and after PLSDA-batch correction of the metagenomic operational taxonomic unit (mOTU) and Kyoto Encyclopedia of Genes and Genomes orthology (KO) blocks; the protein block was analysed without correction. **B**: principal component analysis (PCA) at weeks 0, 8 and 24 for the original and corrected mOTU and KO blocks and the original protein block. Correction reduced the batch-associated separation, although the KO block retained some batch-associated variation at week 24 (Table E.1).

We addressed this technical variation with PLSDA-batch, a supervised approach that removes batch-related variation while maximising the variance explained by treatment (Wang and Lê Cao, 2023). This preservation of treatment-related signal was appropriate for the downstream classification analysis. The protein block showed no detectable batch pattern and was analysed without correction.

We quantified the variation associated with batch and treatment before and after correction (Table E.1). Before correction, batch explained 12.6–26.5% of the main variation in mOTUs and KOs at each week, exceeding the level reached by shuffled labels. After correction, the association was within the shuffled-label range for every block and week except KOs at week 24 (8.4%, compared with 4.8%). Treatment accounted for

**Table E.1. Variation associated with batch and treatment before and after PLSDA-batch correction.** Values are the proportions of variation across the first five principal components (%) associated with each factor in each block and week. Brackets contain the 95th percentile under shuffled labels, and bold values exceed this reference. The protein block was analysed without correction. Because batch groups were inferred from the same data, the batch columns provide a consistency check.

| | | Batch | | Treatment | |
|---|---|---|---|---|---|
| Block | Week | Original | Corrected | Original | Corrected |
| mOTU | 0 | **17.5 (3.7)** | 0.0 (2.8) | 0.5 (3.5) | 1.1 (2.8) |
| mOTU | 8 | **17.0 (3.5)** | 0.0 (2.9) | 3.0 (3.7) | **4.0 (2.8)** |
| mOTU | 24 | **12.6 (3.2)** | 0.4 (2.7) | 0.2 (3.3) | 0.3 (2.7) |
| KO | 0 | **26.5 (6.2)** | 0.1 (5.9) | 0.8 (6.5) | 2.0 (6.1) |
| KO | 8 | **22.6 (6.8)** | 1.1 (8.1) | **11.5 (6.3)** | **16.2 (7.1)** |
| KO | 24 | **16.3 (6.3)** | **8.4 (4.8)** | 0.4 (6.2) | 0.5 (4.6) |

**Table E.2. Sensitivity of BER to batch correction and the block-to-block design weight.** All block.tPLS-DA models used three blocks and three components. Balanced error rate (BER) is the mean (SD) across 10 repeats of three-fold subject-level cross-validation; lower values are better and chance is approximately 0.50. Shuffled-label values are median BERs across 100 permutations, with unadjusted add-one $p$-values. A permutation null was completed only for weight 1. PLSDA-batch used treatment labels from all subjects, including those held out, making the corrected-data BER optimistic.

| Weight | Data | BER | Shuffled labels | $p$ |
|---|---|---|---|---|
| 1 | Uncorrected | 0.465 (0.098) | 0.512 | 0.33 |
| 1 | Corrected | 0.266 (0.065) | 0.521 | 0.03 |
| 0.1 | Uncorrected | 0.438 (0.065) | – | – |
| 0.1 | Corrected | 0.317 (0.090) | – | – |

little of the main variation before or after correction, with the exception of week 8 (KOs: 11.5% before and 16.2% after correction; mOTUs: 4.0% after correction). The correction therefore reduced batch-associated variation while retaining treatment-associated variation. Because the batch groups were defined from the same data, this result provides a consistency check rather than independent evidence of the removal of a technical batch effect.

The effect of correction on classification and its sensitivity to the design weight were assessed by fitting block.tPLS-DA to uncorrected and corrected data with block-to-block weights of 1 and 0.1 (Table E.2). The revised main analysis used weight 1, whereas the submitted analysis used weight 0.1. Correction reduced the BER from 0.465 to 0.266 at weight 1 and from 0.438 to 0.317 at weight 0.1. Differences between the weights were smaller than the variation across cross-validation repeats. The weight-0.1 BERs are descriptive because no permutation null was completed for this sensitivity analysis.

The design weight affected the fitted temporal interpretation more strongly than the held-out error. On the corrected data, the three components used faces 1, 3 and 2 at weight 1, compared with faces 1, 2 and 1 at weight 0.1. Component 1 had a similar direction at both weights (block-wise loading correlations 0.93–0.96), although its highest-ranked features changed. Component 2 shifted from the transient week-8 contrast at weight 1 to the baseline-to-week-24 contrast at weight 0.1. Thus, the weight-0.1 analysis supported the broad Component 1 group contrast, while the specific Component 2 interpretation depended on the design weight.

At both weights, the lower BER after correction was an optimistic estimate of classification performance because PLSDA-batch used treatment labels from all subjects, including those subsequently held out during cross-validation. This procedure introduced information from the test folds into preprocessing.
